# Pyramidal cell types and 5-HT_2A_ receptors are essential for psilocybin’s lasting drug action

**DOI:** 10.1101/2024.11.02.621692

**Authors:** Ling-Xiao Shao, Clara Liao, Pasha A. Davoudian, Neil K. Savalia, Quan Jiang, Cassandra Wojtasiewicz, Diran Tan, Jack D. Nothnagel, Rong-Jian Liu, Samuel C. Woodburn, Olesia M. Bilash, Hail Kim, Alicia Che, Alex C. Kwan

**Affiliations:** Meinig School of Biomedical Engineering, Cornell University, Ithaca, NY, 14853, USA; Department of Psychiatry, Yale University School of Medicine, New Haven, Connecticut, 06511, USA; Interdepartmental Neuroscience Program, Yale University School of Medicine, New Haven, Connecticut, 06511, USA; Medical Scientist Training Program, Yale University School of Medicine, New Haven, Connecticut, 06511, USA; Graduate School of Medical Science and Engineering, KAIST, Daejeon, 34141, Republic of Korea; Department of Psychiatry, Weill Cornell Medicine, New York, NY, 10065, USA

**Author notes:** These authors contributed equally to the work (Ling-Xiao Shao, Clara Liao). Correspondence to Alex Kwan, Ph.D., Room 111 Weill Hall, 526 Campus Road, Ithaca, NY, 14853, United States.

**Keywords:** Psychedelic, serotonin, dendritic spines, structural plasticity, frontal cortex, depression

## Abstract

Psilocybin is a serotonergic psychedelic with therapeutic potential for treating mental illnesses^1–4^. At the cellular level, psychedelics induce structural neural plasticity^5,6^, exemplified by the drug-evoked growth and remodeling of dendritic spines in cortical pyramidal cells^7–9^. A key question is how these cellular modifications map onto cell type-specific circuits to produce psychedelics’ behavioral actions^10^. Here, we use *in vivo* optical imaging, chemogenetic perturbation, and cell type-specific electrophysiology to investigate the impact of psilocybin on the two main types of pyramidal cells in the mouse medial frontal cortex. We find that a single dose of psilocybin increased the density of dendritic spines in both the subcortical-projecting, pyramidal tract (PT) and intratelencephalic (IT) cell types. Behaviorally, silencing the PT neurons eliminates psilocybin’s ability to ameliorate stress-related phenotypes, whereas silencing IT neurons has no detectable effect. In PT neurons only, psilocybin boosts synaptic calcium transients and elevates firing rates acutely after administration. Targeted knockout of 5-HT_2A_ receptors abolishes psilocybin’s effects on stress-related behavior and structural plasticity. Collectively these results identify a pyramidal cell type and the 5-HT_2A_ receptor in the medial frontal cortex as playing essential roles for psilocybin’s long-term drug action.

## Main

Psilocybin is a classic psychedelic that has shown promise as a treatment for psychiatric disorders. Clinical trials demonstrated that one or two sessions of psilocybin-assisted therapy attenuate depression symptoms for many weeks^1–3^. It has been hypothesized that antidepressants may work by forming and strengthening synapses in the prefrontal cortex, which counteracts synaptic dysfunction in depression^11^. Consistent with this framework, recent studies in mice demonstrated that a single dose of psilocybin or related psychedelic drugs leads to sustained increases in the density and size of apical dendritic spines in cortical pyramidal _cells7-9,12,13._

However, neurons are heterogeneous, and it is unclear how psychedelic-evoked neural adaptations manifest in different excitatory cell types. Notably, there are two major, non-overlapping populations of cortical pyramidal cells: pyramidal tract (PT) and intratelencephalic (IT) neurons. PT and IT neurons have distinct cellular properties and participate in different long-range circuits because they send disparate axonal projections to communicate with different brain regions^14–16^ (**Fig. 1a**). PT neurons are subcortical projection neurons that send axons to subcerebral destinations including the thalamus and brainstem, and also to ipsilateral cortex and basal ganglia^16^. By contrast, axons of IT neurons stay within the cerebrum, but can project to both ipsilateral and contralateral cortical and striatal locations. These pyramidal cell types constitute a microcircuit motif that is found in most regions in the neocortex, supporting a range of behavioral functions^17–19^. Impairments of these distinct types of pyramidal cells have been linked to neuropsychiatric disorders^16,20^.

**Fig. 1:**
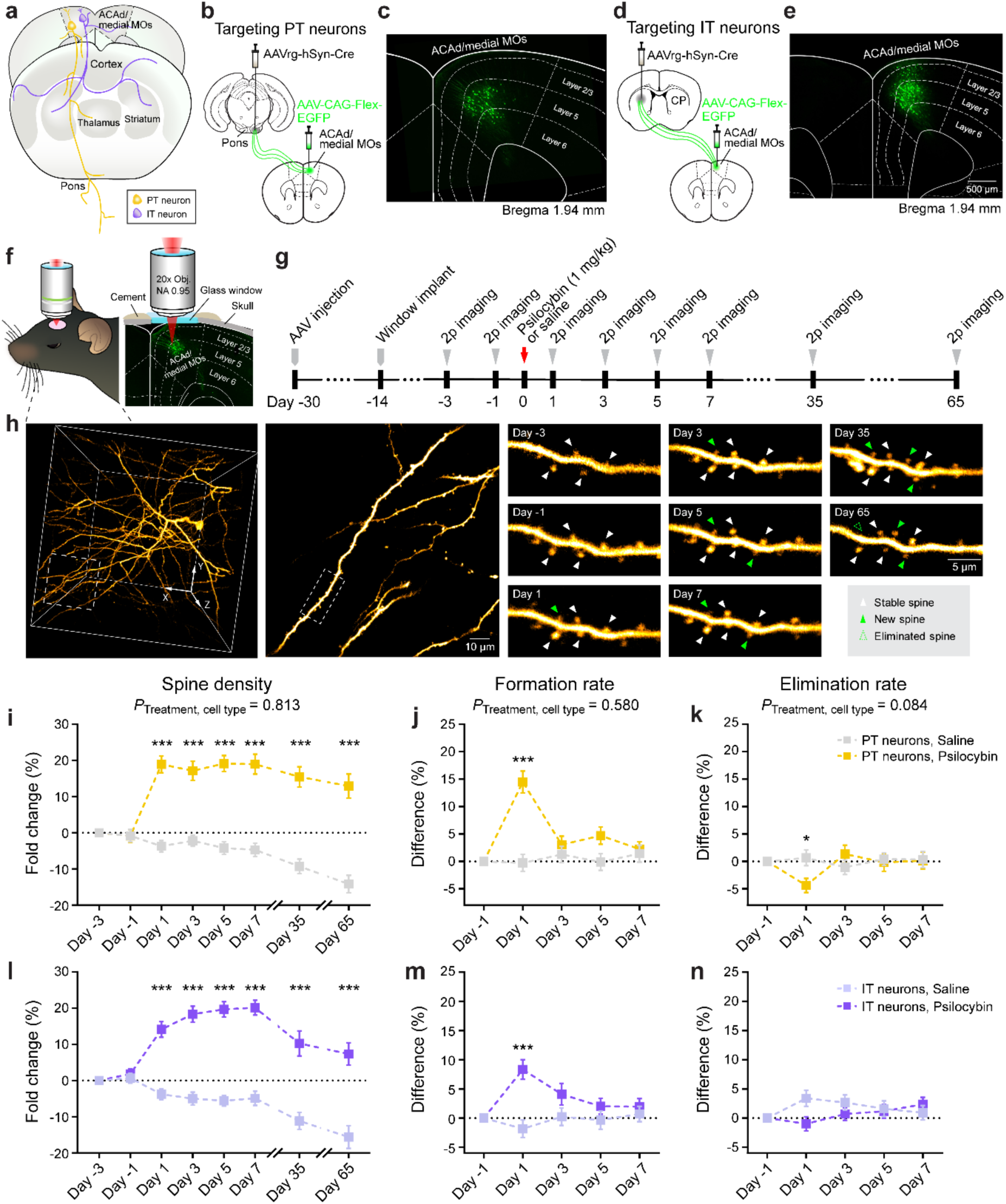
Psilocybin induces structural plasticity in both PT and IT types of frontal cortical pyramidal neurons. **a,** Pyramidal tract (PT) and intratelencephalic (IT) neurons have different long-range projections. **b-c,** Viral strategy to express EGFP selectively in PT neurons in the medial frontal cortex. AAVrg, AAV serotype retrograde. **d-e,** Similar to b-c for IT neurons. CP, caudoputamen. **f-g**, Longitudinal two-photon microscopy. **h,** Example field of view, tracking the same apical tuft dendrites for 65 days after psilocybin. **i,** Density of dendritic spines in the apical tuft of PT neurons after psilocybin (yellow; 1 mg/kg, i.p.) or saline (gray) across days, expressed as fold-change from baseline in first imaging session (day −3). Mean and s.e.m. across dendrites. **j**, Spine formation rate determined by number of new and existing spines in consecutive imaging sessions across two-day interval, expressed as difference from baseline in first interval (day −3 to day −1). **k,** Similar to j for elimination rate. **l-n,** Similar to i-k for IT neurons after psilocybin (purple) or saline (light purple). There was no cell-type difference in psilocybin’s effect on spine density, formation rate, or elimination rate (p-values for interaction effect of treatment × cell type, indicated in plots, mixed effects model). *, p < 0.05. ***, p < 0.001, *post hoc* with Bonferroni correction for multiple comparisons. Sample size *n* values are provided in Methods. Statistical analyses are provided in Supplementary Table 1.

How may PT and IT neurons respond to psilocybin? Classic psychedelics are agonists at serotonin receptors. In response to serotonin, some pyramidal cells elevate spiking activity via 5-HT_2A_ receptors, whereas other pyramidal cells suppress firing via 5-HT_1A_ receptors^21,22^. It was reported that in mouse brain slices, serotonin-evoked firing occurs in pyramidal cells with commissural projections (IT neurons), but not those with corticopontine projections (PT neurons)^23,24^. Transcript expression in the mouse frontal cortex corroborates this view: although PT and IT neurons both express *Htr2a*^25^, there is more *Htr2a* in IT neurons^26^. However, another study performed in anesthetized rats showed that psychedelics can excite midbrain-projecting pyramidal cells, which would constitute PT neurons^27^. Therefore, current literature provides conflicting clues towards how the main pyramidal cell types should contribute to psychedelic drug action.

In this study, we measured the acute and long-term impact of psilocybin on PT and IT neurons in the mouse medial frontal cortex *in vivo*. We found that PT neurons were the pyramidal cell type selectively driven by psilocybin to increase synaptic calcium transients and elevate spiking activity in awake animals. Moreover, although psilocybin evokes structural plasticity in both PT and IT neurons, causal manipulations indicate that frontal cortical PT neurons are needed for psilocybin’s effects in stress-related behavioral assays. Using conditional knockout mice, we found that 5-HT_2A_ receptor is required for psilocybin-evoked structural remodeling in PT neurons. The results thus reveal frontal cortical PT neurons and 5-HT_2A_ receptor as essential components mediating psilocybin’s long-term drug action in the brain.

### Cell-type specificity in the structural plasticity induced by psilocybin

To sparsely express EGFP in PT or IT neurons for dendritic imaging, we injected a low titer of the retrogradely transported AAVretro-hSyn-Cre in the ipsilateral pons or contralateral striatum, and AAV-CAG-FLEX-EGFP in the medial frontal cortex of adult C57BL/6J mice (**Fig. 1b, d; Extended Data Fig. 1**). We focused on the cingulate and premotor portion (ACAd/medial MOs) of the medial frontal cortex, because brain-wide c-Fos mapping indicates the region robustly responds to stress^28^ and psilocybin^29^. Histology confirmed that EGFP-expressing cell bodies of PT neurons were restricted to deep cortical layers, whereas somata of IT neurons were spread across layers 2/3 and 5 (**Fig. 1c, e**), in agreement with the laminar distribution of the cell types^14,16^. We used two-photon microscopy to image through a chronically implanted glass window while the animal was anesthetized. We visualized the same apical tuft dendrites located at 20 – 120 µm below the pial surface over multiple sessions across >2 months (**Fig. 1f–h**). At baseline, PT neurons had lower spine density but higher spine head width than IT neurons (**Extended Data Fig. 2**).

For each of the four cell-type and treatment conditions, we tracked and analyzed 1040–1147 spines from 69–85 dendrites in 8–9 mice of both sexes. For statistical tests, mixed effects models were used, which included random effects terms to account for the nested nature of the data where spines are imaged from the same dendrites or same mouse. Details for sample sizes and statistical tests for all experiments are provided in **Supplementary Table 1**. One dose of psilocybin (1 mg/kg, i.p.) increased spine density in both pyramidal cell types (PT: 19±2% for psilocybin, −4±2% for saline on day 1; IT: 14±2% for psilocybin, −4±1% for saline; main effect of treatment: *P* < 0.001, mixed effects model; **Fig. 1i, l; Extended Data Fig. 3, 4**). The elevated number of dendritic spines remained significant in the last imaging session at 65 days for psilocybin relative to control. For both cell types, the higher spine density was driven by an increase in the rate of spine formation within 1 day after psilocybin (**Fig. 1k, o; Extended Data Fig. 4**), with additionally a smaller decrease in spine elimination rate for PT neurons (**Fig. 1l, p; Extended Data Fig. 4**).

The psilocybin-evoked structural remodeling occurred in mice of both sexes (**Extended Data Fig. 5**). There was no change detected in spine protrusion length (**Extended Data Fig. 6**). Due to the sparse labeling, we could often trace the dendrites back to the cell body. Separately analyzing IT neurons residing in layer 2/3 and layer 5 (**Extended Data Fig. 7**) indicated that laminar position is not the reason for the difference observed across cell type. These results replicate our prior finding^7^ that psilocybin increases spine density in frontal cortical pyramidal cells, while extending the observation window to show that the change persists for >2 months in mice, which occurs for both the PT and IT subpopulations.

### Frontal cortical PT neurons are key for psilocybin’s effect on stress-related behavior

An important question is whether the frontal cortical cell types are relevant for psilocybin’s behavioral effects. To answer this question, we expressed broadly and bilaterally inhibitory DREADD^30^ in PT and IT neurons by injecting AAV-hSyn-DIO-hM4DGi-mCherry in adult *Fezf2-CreER* and *PlexinD1-CreER* mice (**Fig. 2a, b**). These tamoxifen-inducible Cre-driver lines target PT and IT neurons respectively^31^. Control mice were injected with AAV-hSyn-DIO-mCherry. We treated animals with the chemogenetic ligand deschloroclozapine^32^ (DCZ; 0.1 mg/kg, i.p.) 15 minutes before injecting psilocybin (1 mg/kg, i.p.) or saline, thereby silencing the respective subsets of pyramidal cells when the drug is active in the brain.

**Fig. 2:**
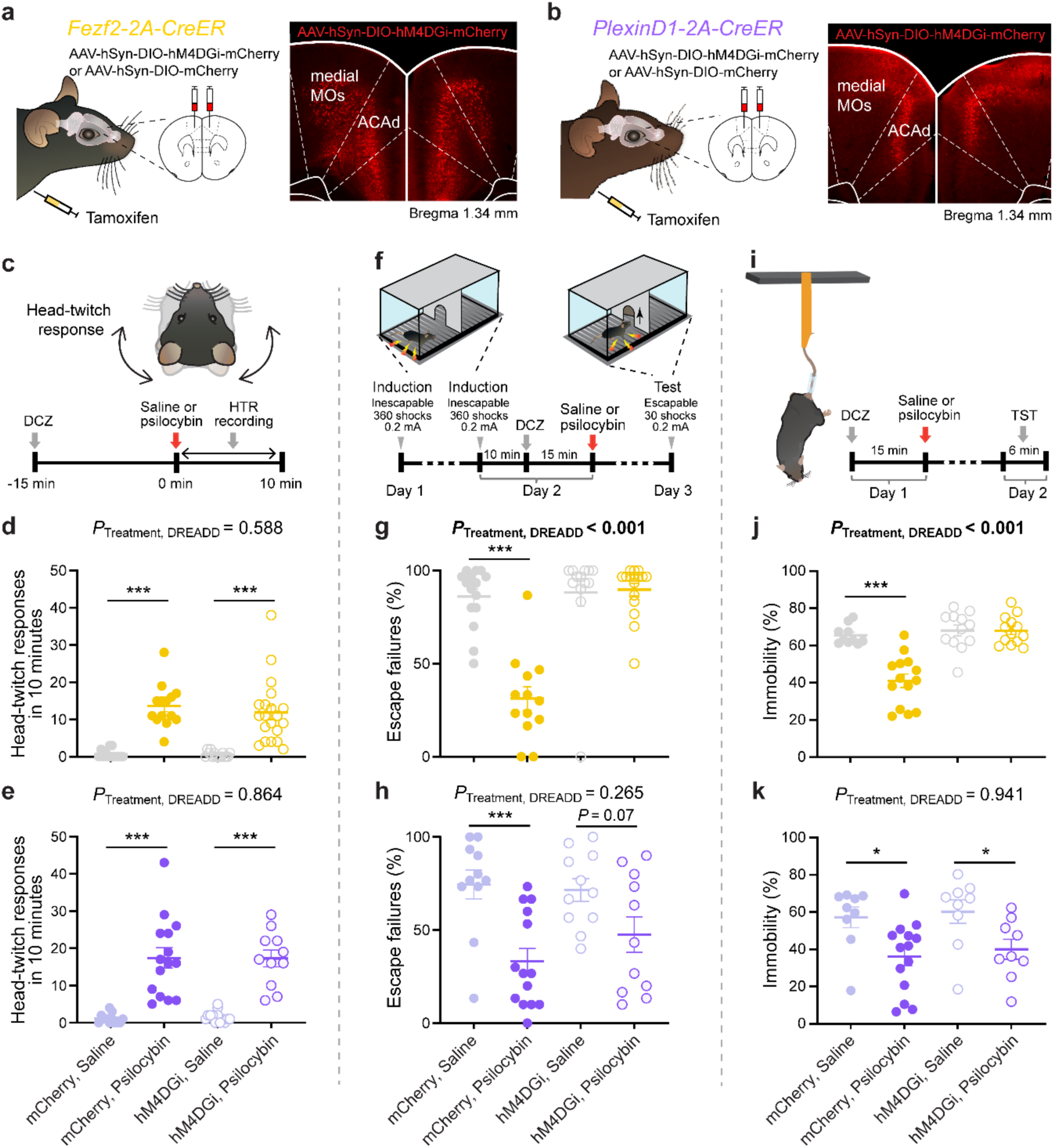
PT neurons are essential for psilocybin’s effects on stress-related behaviors. **a,** Inhibitory chemogenetic receptor (hM4DGi) expressed in PT neurons in the medial frontal cortex of Fezf2-CreER mice. **b,** Similar to a for IT neurons in PlexinD1-CreER mice. **c,** Head-twitch response. **d,** Effect of PT neuron inactivation during psilocybin (1 mg/kg, i.p.) or saline administration. Circle, individual animal. Mean and s.e.m. **e,** Similar to d for IT neurons. **f,** Learned helplessness. **g**, Effect of PT neuron inactivation during psilocybin or saline administration (interaction effect of treatment × DREADD: *P* < 0.001, two-factor ANOVA). Circle, individual animal. Mean and s.e.m. **h,** Similar to g for IT neurons. **i,** Tail suspension test. **j,** Effect of PT neuron inactivation during psilocybin or saline administration on subsequent proportion of time spent immobile (interaction effect of treatment × DREADD: *P* < 0.001, two-factor ANOVA). Circle, individual animal. Mean and s.e.m. **k,** Similar to j for IT neurons. *, p < 0.05. ***, p < 0.001, *post hoc* with Bonferroni correction for multiple comparisons. Sample size *n* values are provided in Methods. Statistical analyses are provided in Supplementary Table 1

We tested four behavioral assays. The head-twitch response is an indicator of hallucinogenic potency of a compound in humans^33^ and occurs nearly immediately after the administration of psilocybin in rodents. Psilocybin induced head twitches in our mice as expected, which was not affected by the DREADD-mediated silencing of frontal cortical PT or IT neurons (n = 11-20 mice in each group; **Fig. 2c-e; Extended Data Fig. 8**). Next, learned helplessness is a preclinical paradigm relevant for modeling depression pathophysiology. Mice were exposed to inescapable footshocks during two induction sessions and subsequently tested for avoidance when faced with escapable footshocks during a test session (**Fig. 2f**). A single dose of psilocybin reduced escape failures, suggesting that drug-treated animals were less affected by the uncontrollable stress (**Fig. 2g, h**). This psilocybin-induced relief of the stress-induced phenotype was abolished if frontal cortical PT neurons were silenced during drug administration (interaction effect of treatment and DREADD: *P* < 0.001, two-factor ANOVA; n = 13-16 mice in each group; **Fig. 2g; Extended Data Fig. 8**). Meanwhile, inactivating IT neurons had no effect (n = 11-14 mice in each group; **Fig. 2h; Extended Data Fig. 8**). The tail suspension test assesses stress-related escape, in which immobility time serves as an indicator of stress-induced escape behavior (**Fig. 2i**). Mice treated with psilocybin 24 hours prior to testing showed a significant reduction in immobility time compared to saline-treated animals, an improvement that was likewise abolished specifically by inactivation of frontal cortical PT neurons (interaction effect of treatment and DREADD: *P* < 0.001, two-factor ANOVA; n = 10-14 mice in each group; **Fig. 2j, k; Extended Data Fig. 8**). Finally, we found that frontal cortical PT neurons are needed for psilocybin-driven facilitation of fear extinction in chronically stressed mice (**Extended Data Fig. 9**). Together the behavioral data indicate that PT neurons in the medial frontal cortex are a key part of the brain’s circuitry for mediating psilocybin’s effect on stress-related behaviors.

### Psilocybin acutely elevates dendritic calcium signaling in PT neurons

What are the early events that initiate the psilocybin-induced structural and behavioral adaptations? Calcium is a second messenger that regulates synaptic plasticity in pyramidal cells^34^. There are different plasticity mechanisms that depend on calcium elevations, both globally in dendritic branches^35^ and locally in dendritic spines^36^. To determine whether calcium in dendritic branches and dendritic spines are involved in psilocybin’s action, we used two-photon microscopy to image the apical dendrites of pyramidal cells in ACAd/medial MOs of awake, head-fixed mice. We focused on the acute phase of psilocybin action, imaging the same fields of view located at 20 – 120 µm below the pial surface for 10 minutes before and within 1 hr after drug injection (**Fig. 3a**). To visualize calcium transients, we expressed the genetically encoded calcium indicator GCaMP6f in PT or IT neurons by injecting AAVretro-hSyn-Cre in the ipsilateral pons or contralateral striatum respectively, and AAV-CAG-FLEX-GCaMP6f in the medial frontal cortex (**Fig. 3b, c**). We used automated procedures^37^ to detect calcium events in regions of interest corresponding to dendritic branches and dendritic spines before and after administering psilocybin (1 mg/kg, i.p.) or saline.

**Fig. 3:**
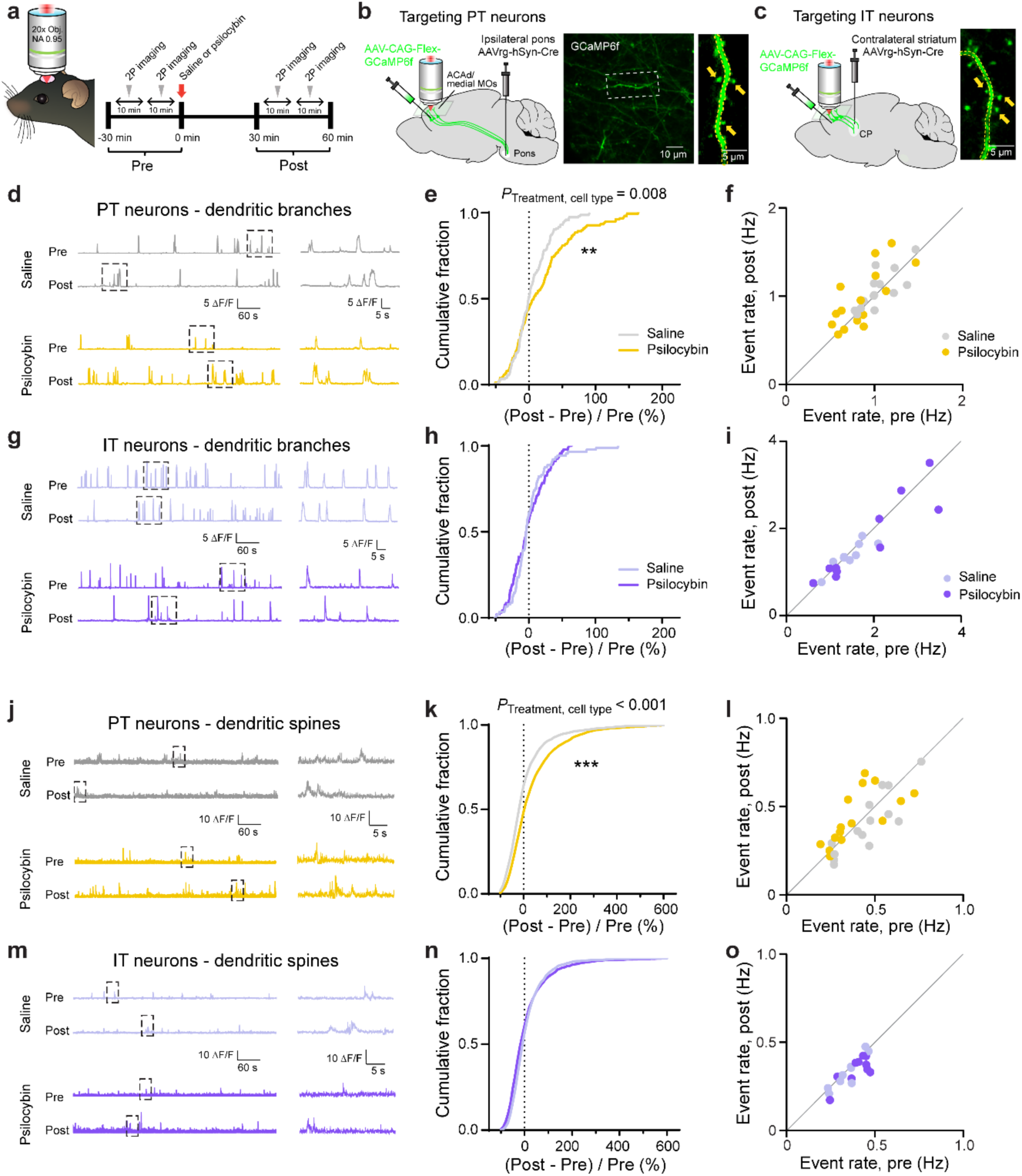
Psilocybin elevates the number of Ca^2+^ transients in dendritic branches and spines of PT neurons. **a,** Two-photon microscopy of spontaneous dendritic calcium signals in awake mice. **b,** Viral strategy to express GCaMP6f selectively in PT neurons in the medial frontal cortex and example *in vivo* image. **c,** Similar to b for IT neurons. **d,** Δ*F*/*F*_0_ from a PT dendritic branch before and after saline or, from a different branch, before and after psilocybin (1 mg/kg, i.p.). Inset (right), magnified view of the boxed area (left). **e,** Fractional change in the rate of calcium events detected in PT dendritic branches after psilocybin (yellow) or saline (gray). **f,** The raw rates of calcium events, averaged across dendritic branches in the same field of view, after psilocybin (yellow) or saline (gray). Each circle is a field of view. **g-i,** Similar to d-f for IT dendritic branches. **j-l,** Similar to d-f for PT dendritic spines. **m-o,** Similar to d-f for IT dendritic spines. **, p < 0.01. ***, p < 0.001, *post hoc* with Bonferroni correction for multiple comparisons. Sample size *n* values are provided in Methods. Statistical analyses are provided in Supplementary Table 1.

For dendritic branches, a single dose of psilocybin increased the rate of spontaneous calcium events in PT neurons (psilocybin: 23±4%; n = 149 branches, 4 mice; saline: 5±2%, n = 140 branches, 4 mice; **Fig. 3d-f; Extended Data Fig. 10, 11**). Conversely, psilocybin did not affect calcium events in dendritic branches of IT neurons (psilocybin: −2±3%; n = 95 branches, 3 mice; saline: 1±3%, n = 90 branches, 3 mice; interaction effect of treatment × cell type: *P* = 0.008, mixed effects model; **Fig. 3g-i; Extended Data Fig. 10, 11**). For dendritic spines, we analyzed fluorescence signals after subtracting contribution from adjoining dendritic branch using a regression procedure^38,39^ to estimate calcium transients arising from subthreshold synaptic activation. Similar to what we saw for dendritic branches, psilocybin elevated the rate of synaptic calcium events in dendritic spines of PT neurons (psilocybin: 68±5%; n = 2637 spines, 4 mice; saline: 37±6%, n = 2307 spines, 4 mice; **Fig. 3j-l; Extended Data Fig. 10, 11**), but not in IT neurons (psilocybin: 20±3%; n = 2198 spines, 3 mice; saline: 16±2%, n = 2237 spines, 3 mice; interaction effect of treatment × cell type: *P* < 0.001, mixed-effects model; **Fig. 3m-o; Extended Data Fig. 10, 11**). These data show that psilocybin preferentially boosts dendritic and synaptic calcium signaling in PT neurons in the medial frontal cortex.

### Psilocybin selectively increases firing in a subset of PT neurons

The heightened dendritic calcium signals are likely due to increased dendritic excitability, which can lead to higher spiking activity in PT neurons. Alternatively, it has been shown that some 5-HT_1A_ receptors localize to the axon initial segment^40^, creating a scenario where dendrites can be excitable while firing remains unchanged or suppressed in PT neurons. To disambiguate these possibilities, we used cell-type specific electrophysiology to record from PT and IT neurons in awake, head-fixed mice. To identify the cell type, we expressed channelrhodopsin (ChR2) in PT or IT neurons by injecting AAV-EF1a-double floxed-hChR2(H134R)-EFYP into the medial frontal cortex of adult *Fezf2-CreER* or *PlexinD1-CreER* mice (**Fig. 4a, b**). We targeted the medial frontal cortex with a high-density Neuropixels probe^41^ (**Fig. 4c**) and isolated single units via spike sorting and quality metrics (**Extended Data Fig. 12**). We recorded for 30 minutes, injected psilocybin (1 mg/kg, i.p.) or saline, and then recorded for another 60 minutes. At the end of each recording session, we performed “opto-tagging” by applying trains of brief laser pulses (473 nm, 20 ms) to identify ChR2-expressing cells. The opto-tagged PT and IT neurons were reliably driven by the photostimulation to spike with short latency (**Fig. 4d-f; Extended Data Fig. 13**).

**Fig. 4:**
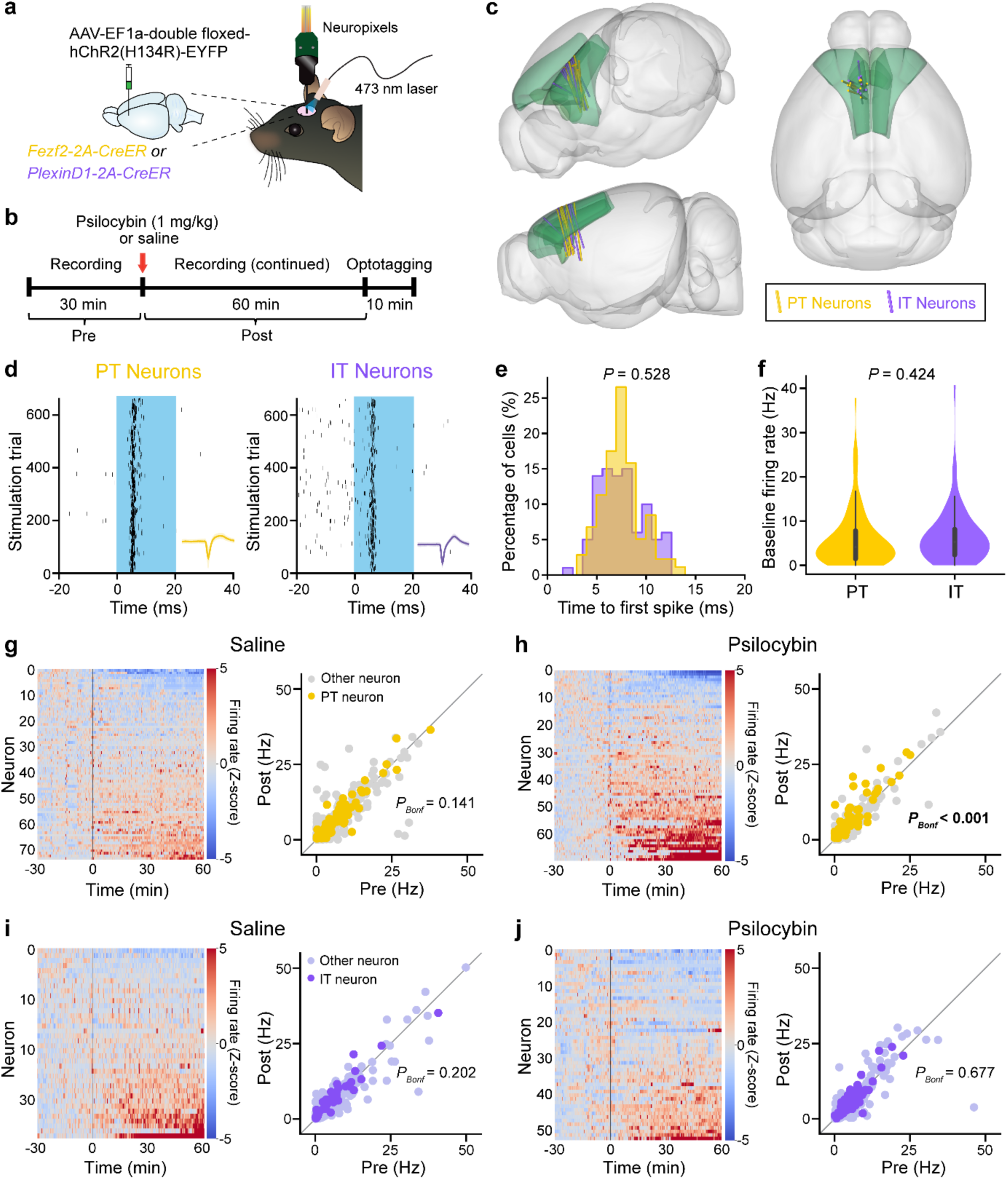
Psilocybin acutely increases firing in a subset of PT neurons *in vivo*. **a,** Neuropixels recording of ChR2-expressing neurons in *Fezf2-CreER* and *PlexinD1-CreER* mice. **b,** Timeline, including pre-drug (−30 – 0 min) and post-drug (0 – 60 min) periods. **c,** Probe tracks recovered from histology and rendered in Allen Mouse Brain Common Coordinate Framework. Green, ACAd and MOs. **d,** Spike raster of tagged neurons. Blue, laser stimulation. Inset, average waveform. **e,** Time of first spike relative to onset of laser for all tagged neurons. Yellow, *Fezf2-CreER*. Purple, *PlexinD1-CreER*. **f,** Mean pre-drug firing rates of all tagged neurons. **g,** Heatmaps showing activity for all tagged neurons in *Fezf2-CreER* mice before and after saline or psilocybin. Firing rate of each neuron was converted to z-score by normalizing based on its pre-drug firing rate. **h,** Mean pre- and post-drug firing rates for all tagged (yellow) and untagged other neurons (gray) in *Fezf2-CreER* mice. Each dot represents one neuron. **i-j,** Similar to g-h for *PlexinD1-CreER* mice. *P_Bonf_*: Bonferroni corrected value. Sample size *n* values are provided in Methods. Statistical analyses are provided in Supplementary Table 1.

A fraction of the opto-tagged PT neurons in *Fezf2-CreER* mice responded vigorously to psilocybin. Specifically, 16% of the PT neurons substantially increased spiking activity, whereas few cells exhibited decrease after psilocybin or change after saline (psilocybin: 14 cells with post-drug mean z-score *Z*>2, 2 cells with *Z*<-2, n = 90 tagged neurons, 5 mice; saline: 2 cell with *Z*>2 and 3 cell with *Z*<-2, n = 104 tagged neurons, 6 mice; **Fig. 4g**). On average, comparing between pre-versus post-drug firing, PT neurons showed significantly higher spike rates after psilocybin (*P* < 0.001, paired *t*-test with Bonferroni correction; **Fig. 4h**). By contrast, there was no notable change in firing activity for IT neurons in *PlexinD1-CreER* mice after psilocybin administration (psilocybin: 2 cells with *Z*>2, 0 cells with *Z*<-2, n = 57 tagged neurons, 5 mice; saline: 2 cells with *Z*>2 and 0 cells with *Z*<-2, n = 38 tagged neurons, 5 mice; *P* = 1.0, paired *t*-test with Bonferroni correction; **Fig. 4i, j**). These results show that psilocybin produces cell type-specific changes in neural dynamics in the medial frontal cortex, highlighted by a set of PT neurons that responded acutely to drug administration by firing vigorously.

### 5-HT2A receptors in medial frontal cortex mediate psilocybin’s alleviating effect on stress-related behaviors

Our results thus far indicate frontal cortical PT neurons as a target for psilocybin. Does the cell type act through 5-HT_2A_ receptors? Current literature provides conflicting data on whether the 5-HT_2A_ receptor is needed^8,42^ or nonessential^43,44^ for the long-term neural and behavioral effects of psychedelics. The discrepancy may stem in part from the use of constitutive knockout animals and antagonist drugs, which can have unwanted effects on neurodevelopment or other receptors. Therefore, here we took a different approach, using a conditional knockout mouse *Htr2a^f/f^* for region- and cell-type-targeted deletion of 5-HT_2A_ receptors in adult animals^45^. We first asked if there are 5-HT_2A_ receptors in frontal cortical excitatory cell types. Analysis of Allen Institute’s single cell sequencing data^46^ revealed abundant *Htr2a* transcripts in a proportion of frontal cortical PT and IT neurons (**Fig. 5a**). Next, we validated Cre-mediated knockout of 5-HT_2A_ receptors in *Htr2a^f/f^* mice. Following injection of AAV-CaMKII-GFP-Cre into the medial frontal cortex, at the transcript level, qPCR confirmed the absence of *Htr2a* transcript in GFP+ cells (control: 2 mice, knockout: 3 mice; **Fig. 5b, c**). At the synaptic level, we performed whole-cell recordings from GFP+ layer 5 pyramidal cells, which did not exhibit 5-HT-evoked increase in sEPSCs (control: 22 cells from 4 mice, knockout: 23 cells from 4 mice; **Fig. 5d**; **Extended Data Fig. 14**), a 5-HT_2A_ receptor-dependent phenomenon^47^.

**Fig. 5:**
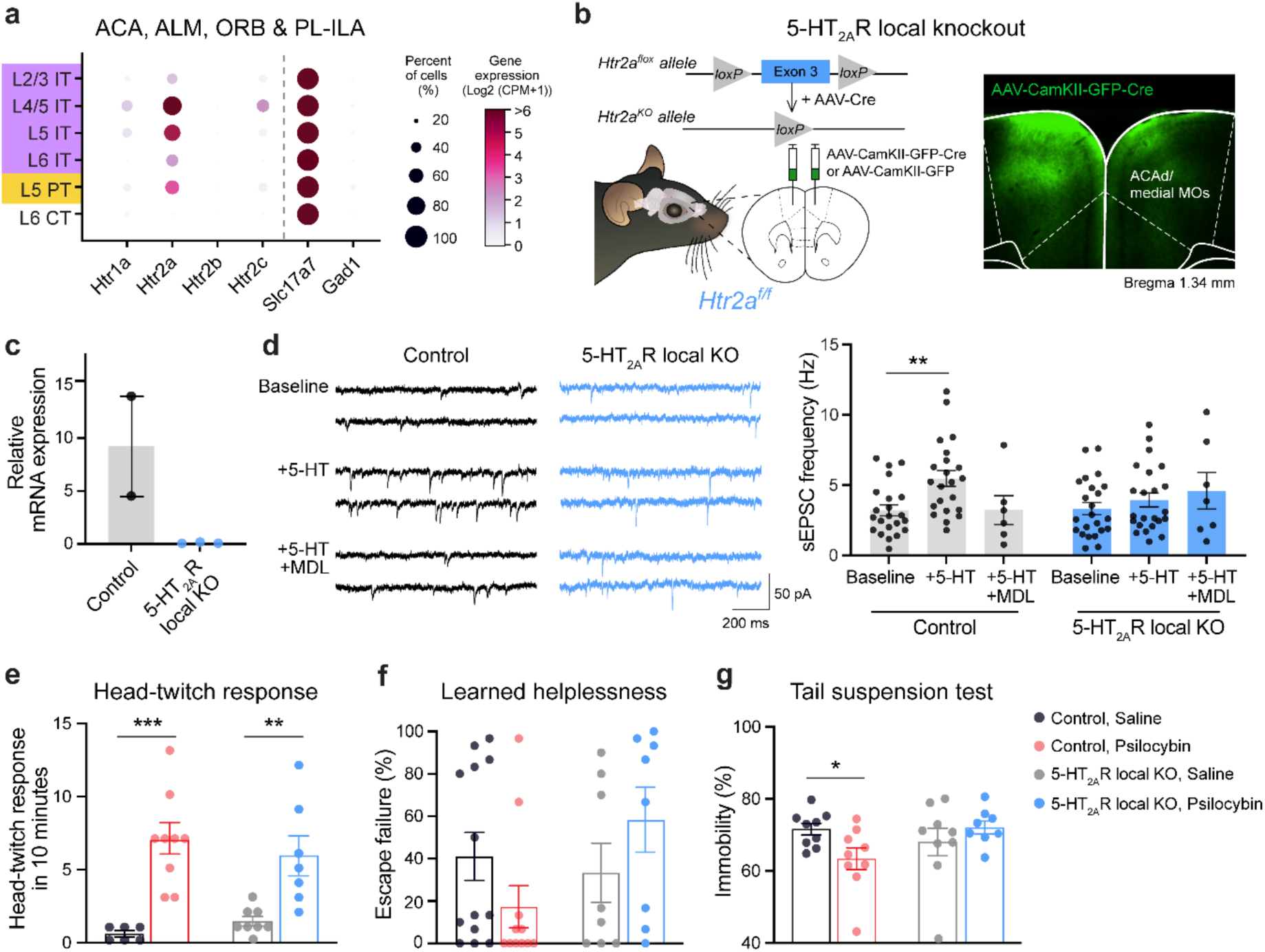
5-HT_2A_ receptors in medial frontal cortex mediate psilocybin’s alleviating effect on stress-related behaviors. **a,** Single-cell transcript counts summed from the mouse ACA, ALM, ORB, and PL-ILA regions, extracted from the SMART-Seq data from Allen Institute. **b,** Viral injection strategy for inducing 5-HT_2A_ receptor knockout or control in GFP+ cells in medial frontal cortex. Inset, post hoc histology. **c,** *Htr2a* mRNA levels in GFP+ cells quantified using qPCR following FACS. Circle, individual animal. **d,** Whole-cell voltage-clamp recordings of spontaneous EPSCs from GFP+ layer 5 pyramidal neurons in baseline, after bath application of 20 µM 5-HT, or after bath application of 20 µM 5-HT and 100 nM MDL100,907. Circle, individual cell. **e,** Head-twitch response measured in control animals after saline (gray) or psilocybin (red) and in animals with bilateral medial frontal cortex-specific 5-HT_2A_ receptor knockout after saline (light gray) or psilocybin (blue). Circle, individual animal. Mean and s.e.m. **f,** Similar to e for learned helplessness. **g,** Similar to e for tail suspension. *, p < 0.05. **, p < 0.01. ***, p < 0.001. Sample size *n* values are provided in Methods. Statistical analyses are provided in Supplementary Table 1.

Leveraging the *Htr2a^f/f^*mice, we asked if the 5-HT_2A_ receptor in frontal cortical neurons are needed for psilocybin’s effects in the same set of behaviors tested for **Fig. 2**. We injected either AAV-hSyn-Cre-P2A-Tomato or AAV-hSyn-EGFP bilaterally and broadly in the medial frontal cortex of *Htr2a^f/f^* mice. Animals with the localized knockout of 5-HT_2A_ receptors exhibited the same amount of psilocybin-evoked head-twitch response as controls (n = 6-9 mice in each group; **Fig. 5e**). The lack of dependence on 5-HT_2A_ receptor for the psilocybin-evoked head-twitch response was specific to local manipulation in the medial frontal cortex, because *CaMKII^Cre^*;*Htr2a^f/f^* mice with constitutive and more widespread receptor knockout had markedly fewer head-twitch response than control animals after psilocybin administration (**Extended Data Fig. 15**). Notably, the region-specific 5-HT_2A_ receptor knockout was sufficient to render psilocybin ineffective for ameliorating the stress-related phenotypes in learned helplessness (n = 8-13 mice in each group; **Fig. 5f**) and tail suspension test (n = 8-9 mice in each group; **Fig. 5g**). We note the caveat that although results from head-twitch response and tail suspension test were clearly interpretable, the response of control animals to psilocybin in learned helplessness did not reach statistical significance, likely due to floor effect from the low baseline rate of escape failures in this *Htr2a^f/f^* strain. Collectively, the data show the importance of 5-HT_2A_ receptors in the medial frontal cortex for psilocybin’s ameliorative effects on stress-related behavior.

### 5-HT2A receptor is required for psilocybin-induced structural plasticity in PT neurons

Is the 5-HT_2A_ receptor needed for psilocybin-evoked dendritic remodeling? To answer this question, we performed targeted knockout of 5-HT_2A_ receptors by injecting low titer of AAVretro-hSyn-Cre into the ipsilateral pons and AAV-CAG-FLEX-EGFP into the medial frontal cortex of *Htr2a^f/f^* (**Fig. 6a**). In this viral strategy, the Cre recombinase was needed for dual purposes to express EGFP for visualization and to mediate knockout, therefore the control animals need the same viruses, which are injected into wild type C57BL/6J mice. We used two-photon microscopy to image the same apical tuft dendrites for 4 sessions including before and after treatment with psilocybin (1 mg/kg, i.p.) or saline (**Fig. 6b, c**). For each condition (genotype and psilocybin or saline), we tracked and analyzed 445–1008 spines from 31–68 dendrites in 5–7 mice of both sexes. In agreement with our earlier findings, the frontal cortical PT neurons in control animals exhibited increased spine density following a single dose of psilocybin (spine density: 15±2% for psilocybin, −2±2% for saline on day 1). By contrast, the cell type-targeted 5-HT_2A_ receptor knockout abolished psilocybin’s effects (spine density: −1±2% for psilocybin, 1±2% for saline on day 1; interaction effect of treatment × genotype: *P* = 0.01 for spine density, mixed effects model; **Fig. 6d-f; Extended Data Fig. 16-17**). *In vivo* two-photon microscopy has spatial resolution close to the limit needed for measuring spine size, motivating us to perform post hoc confocal microscopy in fixed tissues from the same animals to determine psilocybin’s impact on spine morphology (n = 68-136 dendrites from 3-7 mice in each group; **Fig. 6g**). Extracted from day 3 after psilocybin dosing, the confocal data showed a psilocybin-evoked increase of spine head width in apical tufts of frontal cortical PT neurons (0.53±0.01 μm for psilocybin, 0.50±0.01 μm for saline), an effect that was absent when 5-HT_2A_ receptors were selectively deleted (0.50±0.01 μm for psilocybin, 0.51±0.01 μm for saline; interaction effect of treatment × genotype: *P* = 0.025, two-factor ANOVA; **Fig. 6h-i**). These data strongly point to the necessity of 5-HT_2A_ receptors for psilocybin-induced structural neural plasticity.

**Fig. 6:**
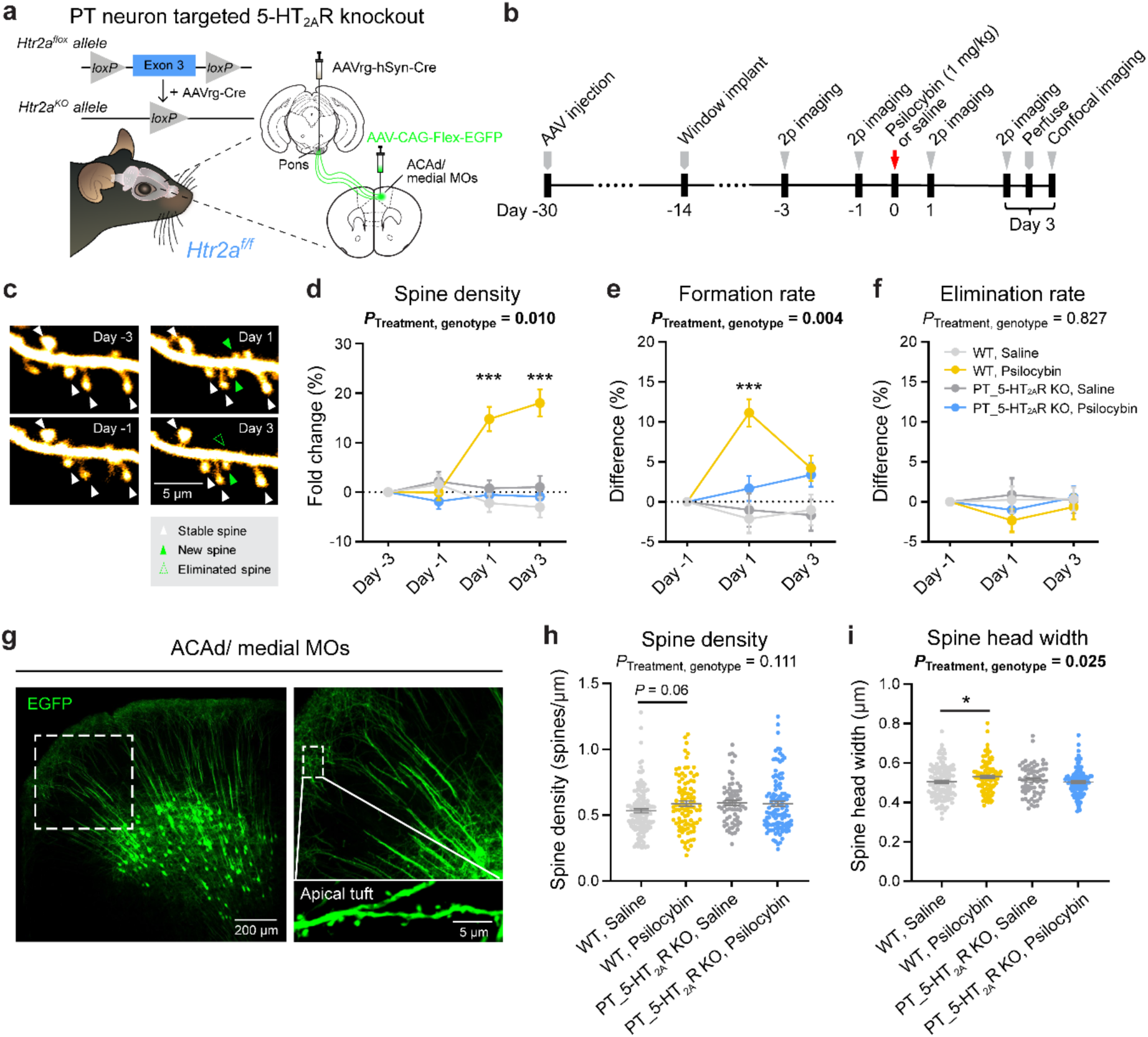
5-HT_2A_ receptor is required for psilocybin-induced structural plasticity in PT neurons. **a,** Viral injection strategy for inducing conditional 5-HT_2A_ receptor knockout and GFP expression for imaging in frontal cortical PT neurons. **b,** Longitudinal two-photon microscopy followed by confocal imaging. **c,** Example field of view, tracking the same apical tuft dendrites before and after psilocybin. **d,** Density of dendritic spines in the apical tuft of PT neurons across days, expressed as fold-change from baseline in first imaging session (day −3), in wild type mice after saline (light gray) or psilocybin (yellow) and in mice with PT neuron-targeted 5-HT_2A_ receptor knockout after saline (gray) or psilocybin (blue). Mean and s.e.m. across dendrites. *Post hoc* test compared WT:saline and WT:psilocybin groups. **e,** Spine formation rate determined by number of new and existing spines in consecutive imaging sessions across two-day interval, expressed as difference from baseline in first interval (day - 3 to day −1. Interaction effect of treatment × cell type: *P* = 0.004, mixed effects model). **f,** Similar to e for elimination rate. **g,** Example field of view imaging apical tufts using confocal microscopy. **h,** Density of dendritic spines in the apical tuft of PT neurons in wild type mice after saline (light gray) or psilocybin (yellow) and in mice with PT neuron-targeted 5-HT_2A_ receptor knockout after saline (gray) or psilocybin (blue). Circle, individual dendritic segment. Mean and s.e.m. **i,** Similar to h for spine head width (interaction effect of treatment × genotype: *P* = 0.025, two-factor ANOVA). *, p < 0.05. ***, p < 0.001, *post hoc* with Bonferroni correction for multiple comparisons. Sample size *n* values are provided in Methods. Statistical analyses are provided in Supplementary Table 1.

## Discussion

We demonstrate that psilocybin’s long-term behavioral effects are dissociated at the level of pyramidal cell types in the frontal cortex. The cell-type specific dissociation may be a mechanism leveraged by novel psychedelic analogs to isolate therapeutic effects from hallucinogenic action^12,48,49^. A key finding is that frontal cortical PT neurons are essential for psilocybin’s beneficial effects in stress-related phenotypes. The consequence for the structural plasticity in frontal cortical IT neurons is unclear; it may be an epiphenomenon, or the IT cell type may mediate other psilocybin-induced behavioral changes that were not tested in this study.

Our results emphasize the importance of 5-HT_2A_ receptors for psilocybin’s long-term effects. However, given that many PT and IT neurons in the frontal cortex have abundant *Htr2a* transcripts, the expression profile cannot fully explain why PT neurons respond preferentially to psilocybin. It is plausible that under *in vivo* conditions, circuit mechanisms steer psilocybin’s action to favor PT neurons. For instance, psilocybin may heighten activity of certain long-range axonal inputs with biased connectivity to frontal cortical PT neurons, such as those from contralateral medial frontal cortex^50^ and ventromedial thalamus^51^. Another possibility is that psilocybin may cause disinhibition by suppressing specific GABAergic neurons, such as deep-lying somatostatin-expressing interneurons that preferentially inhibit PT neurons^52,53^. Receptor and circuit mechanisms are not mutually exclusive and their relative contributions to psilocybin’s impact on frontal cortical neural dynamics should be determined in future studies.

A hallmark of psychedelics is their ability to alter conscious perception. Layer 5 pyramidal cells, including specifically the PT neuron subpopulation, have been implicated in the transition from anesthesia to wakefulness^54,55^. In the medial frontal cortex, PT neurons represent the subcortical output pathway, sending axons to ipsilateral thalamus and other deep-lying brain regions. There is growing interest to develop new treatments for depression that pair antidepressants with other approaches, such as electroconvulsive or transcranial magnetic stimulation^56^, with a goal to augment neural plasticity and enhance therapeutic outcome. This study delineates the cell types and receptors that underpin psychedelic action, highlighting the neural circuits that may be promising targets for neuromodulation and precision treatment.

## Acknowledgments

We thank Liyuan Sun for help with analyzing the slice electrophysiology data. Psilocybin was generously provided by Usona Institute’s Investigational Drug & Material Supply Program; the Usona Institute IDMSP is supported by Alexander Sherwood, Robert Kargbo, and Kristi Kaylo in Madison, WI. This work was supported by NIH grants R01MH121848, R01MH128217, U01NS128660, One Mind – COMPASS Rising Star Award (A.C.K.); NIH training grants T32GM007205 (P.A.D. and N.K.S.), T32NS041228 (C.L.); NIH fellowships F30DA059437 (P.A.D.), F30MH129085 (N.K.S.); Source Research Foundation student grant (P.A.D.); NIH instrumentation grants S10RR025502 and S10OD032251 (Cornell Biotechnology Resource Center Imaging Facility); NIH grants R00NS114166, R01NS133434, and R01DA059378 (A.C.); State of Connecticut, Department of Mental Health and Addiction Services (A.C. and R.L.).

## Contributions

L.X.S., C.L, and A.C.K planned the study. L.X.S. and C.L. conducted and analyzed the imaging and behavioral experiments. P.A.D. and Q.J. conducted and analyzed the electrophysiological experiments. N.K.S. and O.M.B. analyzed the dendritic calcium imaging data. R.J.L and A.C conducted slice electrophysiology experiments. Q.J. assisted in animal surgery. Q.J., D.T., C.W., and J.D.N. assisted in behavioral experiments and histology. S.C.W. and C.W. conducted pilot studies to validate the protocols for the behavioral assays. H.K generated and provided the *Htr2a^f/f^* mice. L.X.S., C.L., and A.C.K. drafted the manuscript. All authors reviewed the manuscript before submission.

## Data availability

Data and code associated with the study will be available on https://github.com/Kwan-Lab.

## Methods

### Animals

Wild-type C57BL/6J (Stock No. 000664), *Fezf2-2A-CreER^1^* (B6;129S4-Fezf2^tm1.1(cre/ERT2)Zjh^/J, Stock No. 036296), *PlexinD1-2A-CreER^1^* (B6;129S4-*Plxnd1^tm^*^1.1^*^(cre/ERT^*^2^*^)Zjh/J^*, Stock No. 036296), *Thy1^GFP^* line M^2^ (Tg(Thy1-EGFP)MJrs/J, Stock No. 007788), and *CaMKIIa^Cre^* (B6.Cg-Tg(Camk2a-cre)T29-1Stl/J, Stock No. 005359) mice were from Jackson Laboratory and bred in our animal facility. *Htr2a^f/f^* mice were described in a previous study^3^ and bred in our animal facility. For behavioral and electrophysiological studies involving PT and IT neurons, 5- to 8-week-old homozygous *Fezf2-2A-CreER* and *PlexinD1-2A-CreER* mice were used for viral injection, then tested 2 weeks later. For two-photon imaging studies, 5 to 7-week-old C57BL/6J or homozygous *Htr2a^f/f^* mice were used for viral injection, then implanted with a glass window and imaged 2-3 weeks later. For validation and behavioral studies involving the *Htr2a^f/f^*mice, 5 to 8-week-old homozygous *Htr2a^f/f^* mice or littermate controls were used for viral injection, then tested 3 weeks later. Animals were housed in groups with 2 – 5 mice per cage in a temperature-controlled room, operating on a normal 12 hr light - 12 hr dark cycle (8:00 AM to 8:00 PM for light). Food and water were available ad libitum. Animals were randomly assigned to different experimental groups. Animal care and experimental procedures were approved by the Institutional Animal Care & Use Committee (IACUC) at Cornell University and Yale University.

### Viruses

AAV1-pCAG-FLEX-EGFP-WPRE (Catalog #51502), AAVretro-hSyn-Cre-WPRE-hGH (Catalog #105553), AAV1-CAG-Flex-GCaMP6f-WPRE-SV40 (Catalog #100835), AAV1-hSyn-DIO-hM4D(Gi)-mCherry (Catalog #44362), AAV1-hSyn-DIO-mCherry (Catalog #50459), AAV1-EF1a-double floxed-hChR2(H134R)-EYFP-WPRE-HGHpA (Catalog #20298), AAV9-CaMKII-HI-GFP-Cre.WPRE.SV40 (Catalog #105551), AAV8-CaMKIIa-EGFP (Catalog #50469), AAV9-hSyn-Cre-P2A-Tomato (Catalog #107738), and AAV9-hSyn-EGFP (Catalog #50465) were purchased from Addgene. AAVretro is an AAV designed for efficient retrograde transport^4^. All viruses had titers ≥ 7×10^12^ vg/mL. The viruses were stored at −80°C. Before stereotaxic injection, they were taken out of the - 80°C freezer, thawed on ice, and diluted to the corresponding titer for injection.

### Surgery

Prior to surgery, each mouse was injected with dexamethasone (3 mg/kg, i.m.; DexaJect, #002459, Henry Schein Animal Health) and carprofen (5 mg/kg, s.c.; #024751, Henry Schein Animal Health) for anti-inflammatory and analgesic purposes. At the start of surgery, anesthesia was induced with 2 – 3% isoflurane and the mouse was affixed in a stereotaxic apparatus (Model 900, David Kopf Instruments). Anesthesia was maintained with 1 – 1.5% isoflurane. Body temperature was maintained at 38°C using a far-infrared warming pad (#RT-0515, Kent Scientific). Petrolatum ophthalmic ointment (#IS4398, Dechra) was applied to cover the eyes. The hair on the head was shaved. The scalp was disinfected by wiping with ethanol pads and povidone-iodine. Small burr holes were made above the targeted brain regions using a handheld dental drill (#HP4-917, Foredom). Adeno-associated virus (AAV) was delivered intracranially into the brain by inserting a borosilicate glass capillary and using an injector (Nanoject II Auto-Nanoliter Injector, Drummond Scientific). Injections were done for the various experiments using different viruses and volumes, as specified in the paragraphs below, using 4.6 nL pulses with 20 s interval between each pulse. To reduce backflow of the virus, we waited 5-10 min after completing an injection at one site before retracting the pipette to move on to the next site. For the medial frontal cortex and striatum, the stereotaxic apparatus was positioned at four sites corresponding to four vertices of a 0.2 mm-wide square centered at the coordinates mentioned below. Throughout the procedure, the brain surface was kept moist with artificial cerebrospinal fluid (aCSF; in mM: 135 NaCl, 5 HEPES, 5 KCl, 1.8 CaCl2, 1 MgCl2; pH: 7.3). After injections, the craniotomies were covered with silicone elastomer (#0318, Smooth-On, Inc.), and the skin was sutured (#1265B, Surgical Specialties Corporation). At the end of surgery, animal was given carprofen (5 mg/kg, s.c.) immediately and then again once on each of the following 3 days.

For two-photon imaging of dendritic structure, to target PT neurons, 110.4 nL of AAVretro-hSyn-Cre-WPRE-hGH (1:100 diluted in phosphate-buffered saline (PBS; #P4417, Sigma-Aldrich)) was injected into the pons (anteroposterior (AP): −3.4 mm, mediolateral (ML): −0.7 mm, dorsoventral (DV): −5.2 mm; relative to bregma, same applies below, unless otherwise specified) and 92 nL of AAV1-pCAG-FLEX-EGFP-WPRE (1:20 diluted in PBS) was injected into the ACAd and medial MOs subregion of medial frontal cortex (AP: 1.5 mm, ML: −0.4 mm, DV: −1.0 mm) of a C57BL/6J or a *Htr2a^f/f^* mouse. To target IT neurons, 101.2 nL of AAVretro-hSyn-Cre-WPRE-hGH (1:100 diluted in PBS) was injected into the contralateral striatum (AP: 0.6 mm, ML: 2.2 mm, DV: −2.8 mm) and 92 nL of AAV1-pCAG-FLEX-EGFP-WPRE (1:20 diluted in PBS) was injected into the medial frontal cortex of a C57BL/6J mouse. After 2-3 weeks, the mouse underwent a second procedure, with the same pre- and post-operative care, to implant a glass window for imaging. An incision was made to remove skin above the skull, and the skull was cleaned to remove connective tissues. A dental drill was used to make a ∼3-mm-diameter circular craniotomy above the previously targeted location at the medial frontal cortex. aCSF was used to immerse the exposed dura in the craniotomy. A two-layer glass window was made bonding two round coverslips (3 mm diameter, 0.15 mm thickness; #640720, Warner Instruments) via ultraviolet light-curing optical adhesive (#NOA 61, Norland Products) using an ultraviolet illuminator (#2182210, Loctite). The glass window was placed over the craniotomy and, while maintaining a slight pressure, super glue adhesive (Henkel Loctite 454) was carefully used to secure the window to the surrounding skull. A stainless steel headplate (eMachineShop; design available at https://github.com/Kwan-Lab/behavioral-rigs) was secured on the skull and centered on the glass window using a quick adhesive cement system (Metabond, Parkell). Mouse would recover for at least 10 days after the window implant prior to imaging experiments.

For two-photon imaging of dendritic calcium transients, to target PT neurons, 110.4 nL of AAVretro-hSyn-Cre-WPRE-hGH (1:10 diluted in PBS) was injected into the pons, and 92 nL of AAV1-CAG-Flex-GCaMP6f-WPRE-SV40 (1:10 diluted in PBS) was injected into medial frontal cortex of a C57BL/6J mouse. To target IT neurons, 110.4 nL of AAVretro-hSyn-Cre-WPRE-hGH (1:10 diluted in PBS) was injected into contralateral striatum and 92 nL of AAV1-CAG-Flex-GCaMP6f-WPRE-SV40 (1:10 diluted in PBS) was injected into medial frontal cortex of a C57BL/6J mouse. For two-photon imaging of dendritic structure in *Thy1^GFP^* mice, 345 nL of AAV9-hSyn-Cre-P2A-Tomato (1:100 diluted in PBS) was injected into the medial frontal cortex of *Thy1^GFP^*;*Htr2a^f/f^* or *Thy1^GFP^*;*Htr2a^WT/WT^* mice. The surgical procedures were the same as above.

For chemogenetic experiments, 276 nL of AAV1-hSyn-DIO-hM4D(Gi)-mCherry or the control virus AAV1-hSyn-DIO-mCherry was injected into the medial frontal cortex bilaterally (AP: 1.5 mm, ML: 0.4 and −0.4 mm, DV: −1.0 and −1.2 mm) of *Fezf2-2A-CreER* mice to target PT neurons and *PlexinD1-2A-CreER* mice to target IT neurons. For *in vivo* electrophysiology, 276 nL of AAV1-EF1a-double floxed-hChR2(H134R)-EFYP was injected into the medial frontal cortex unilaterally (AP: 1.5 mm, ML: 0.4 mm, DV: −1.0 and −1.2 mm) of *Fezf2-CreER* and *PlexinD1-CreER* mice to target PT and IT neurons, respectively. After 2 weeks, the mouse would undergo a second procedure. An incision was made to remove the skin and the periosteum was cleared. A dental drill was used to make a 0.9 mm craniotomy and a 0.86 mm self-tapping bone screw (#19010-10, Fine Science Tools) was placed through the skull bone into the cerebellum to act as a ground screw and provide further structural support for head-fixation. A custom stainless steel headplate was affixed on the skull using a quick adhesive cement system. The mouse would recover for at least one-week post-surgery prior to commencement of electrophysiological experiments. In both cases, injection of AAVs were made prior to administration of tamoxifen.

For validation of the *Htr2a^f/f^* mouse line, homozygous *Htr2a^f/f^* animals were bilaterally injected with AAV9-CaMKII-HI-GFP-Cre.WPRE.SV40 (441.6 nL, 1:10 diluted in PBS) or AAV8-CaMKIIa-EGFP (441.6 nL, 1:10 diluted in PBS) in the medial frontal cortex (AP: 1.5 mm, ML: +/-0.4 mm, DV: −0.4, −0.6, and −1.2 mm, relative to dura). Incised skin was sutured. Animals would recover for at least 3 weeks prior to sacrifice for transcript or slice electrophysiology experiments. For behavioral experiments involving *Htr2a^f/f^*mice, homozygous *Htr2a^f/f^* animals were bilaterally injected with AAV9-hSyn-Cre-P2A-Tomato (690 nL, 1:50 diluted in PBS) or AAV9-hSyn-EGFP (690 nL, 1:50 diluted in PBS) in the medial frontal cortex (AP: 1.5 mm, ML: −0.4 mm, DV: −0.4, −0.6, and −1.2 mm, relative to dura). Incised skin was sutured. Animals would recover for at least 3 weeks prior to behavioral experiments.

### Tamoxifen

Tamoxifen was used for inducible Cre-dependent gene expression in *Fezf2-CreER* and *PlexinD1-CreER* mice. Tamoxifen (#T5648, Sigma-Aldrich) was dissolved in corn oil (#C8267, Sigma-Aldrich) at concentration of 20 mg/mL in an ultrasonic bath at 37°C for 1 – 4 hr. The solution was then aliquoted into 1 mL tubes, wrapped with aluminum foil, and stored at −20°C. For injections, the tamoxifen aliquots were thawed at 4°C. Each animal was weighed and received tamoxifen (75 mg/kg, i.p.) once every 24 hours for 5 consecutive days. Experiments involving inducible Cre expression were conducted at least 2 weeks after the last dose of tamoxifen to allow time for viral-mediated expression.

### Histology

Histology was performed to determine the accuracy of injection locations and assess transgene expression. For two-photon imaging and behavioral studies, after completion of experiments, mice were perfused with PBS, followed by paraformaldehyde solution (PFA, 4% (v/v) in PBS). The brains were extracted and further fixed in 4% PFA at 4°C for 12 – 24 hr. Subsequently, 40-µm-thick coronal sections were obtained using a vibratome (#VT1000S, Leica) and mounted on slides with glass coverslips. Sections were imaged using a wide-field fluorescence microscope (BZ-X810, Keyence). For electrophysiology, the coronal sections were prepared similarly, mounted on slides using Vectashield containing DAPI (#H-1200-10, Vector Laboratories) and imaged. To locate the Neuropixels probe, we used SHARP-TRACK^5^ to align the images of the coronal sections including the DiI tracks with the standardized Allen Common Coordinate Framework 6^6^. Reconstructed probe tracks were visualized within the Allen Common Coordinate Framework using Brainrender^7^.

### Two-photon imaging

Two-photon imaging experiments were performed using a Movable Objective Microscope (MOM, Sutter Instrument) equipped with a resonant-galvo scanner (Rapid Multi Region Scanner, Vidrio Technologies) and a water-immersion 20X objective (XLUMPLFLN, 20x/0.95 N.A., Olympus). ScanImage 2020 software^8^ was used to control the microscope for image acquisition. To visualize GFP or GCaMP6f-expressing dendrites, a tunable Ti:Sapphire femtosecond laser (Chameleon Ultra II, Coherent) was used as the excitation source. The excitation wavelength was set at 920 nm, and emission was collected behind a 475 – 550 nm bandpass filter for fluorescence from GFP or GCaMP6f. The laser power measured at the objective was typically ≤40 mW and varied depending on the imaging depth. When imaging of the same field of view across days, the laser power was kept the same in each imaging session.

For structural imaging of dendrites, in each imaging session, the mouse was head fixed and anesthetized with 1% isoflurane through a nose cone. Body temperature was maintained at 37.4°C via a heating pad system (#40-90-8D, FHC) with feedback control from a rectal thermistor probe. Each imaging session lasted 0.5 – 1.5 hr. To target the ACAd and medial MOs subregion of the medial prefrontal cortex, we imaged within 400 µm of the midline as determined by first visualizing the sagittal sinus in bright-field imaging. To target apical tuft dendrites, we first imaged 0 – 200 µm below the pial surface to identify the apical tuft dendrites and apical trunk, and then select apical tuft dendrites located between 20 – 120 µm below the pial surface for longitudinal imaging. Multiple different fields of view were imaged in the same mouse. For each field of view, 10 – 40-µm-thick z-stacks were collected with 1 µm steps using 15 Hz bidirectional scanning at 1024 × 1024 pixels with a resolution of 0.11 µm per pixel. Each mouse was imaged at the same fields of view on day −3, −1, 1, 3, 5, 7, 35 and 65 relative to the day of drug administration. On the day of treatment (day 0), no imaging was performed, and the mouse was injected while awake with either psilocybin (1 mg/kg, i.p.; prepared from working solution, which was made fresh monthly from powder; Usona Institute) or saline (10 mL/kg, i.p.). After injection, the mouse was placed in a clean cage, and head twitches were visually inspected for 10 min before returning the mice to their home cage. At the end of imaging session, for the purpose of reconstructing the apical dendritic trees, a z-stack was acquired between 0 – 900 µm below the dura with 2 µm steps. For structural imaging of dendrites, 148 dendrites from 17 C57BL/6J mice were imaged for psilocybin (8 males including 5 for PT and 3 for IT neurons; 9 females including 4 for PT and 5 for IT neurons), and 154 dendrites from 16 C57BL/6J mice were imaged for saline (7 males including 4 for PT and 3 for IT neurons; 9 females including 4 for PT and 5 for IT neurons). For structural imaging to test effects of 5-HT_2A_ receptor knockout on dendrites, 117 dendrites from 11 mice were imaged for psilocybin (6 *Htr2a^f/f^* mice; 5 C57BL/6J mice), and 80 dendrites from 12 mice were imaged for saline (5 *Htr2a^f/f^* mice; 7 C57BL/6J mice). For structural imaging of *Thy1^GFP^* mice, 38 dendrites from 2 *Thy1^GFP^*;*Htr2a^WT/WT^* mice were imaged for psilocybin, 49 dendrites from 5 *Thy1^GFP^*;*Htr2a^f/f^*mice were imaged for psilocybin, and 98 dendrites from 3 *Thy1^GFP^*;*Htr2a^f/f^*mice were imaged for saline.

For calcium imaging of dendrites, the mouse was habituated to head–fixation in an acrylic tube under the microscope for 3–4 days, with increasing durations each day, before the day of data collection. To examine the acute effects of psilocybin, we imaged 2 fields of view, each for 10 min to obtain pre-treatment baseline data. Imaging was then paused to inject psilocybin (1 mg/kg, i.p.) or saline (10 mL/kg, i.p.). At 30 min after injection, we imaged those same 2 fields of view again, each for 10 min to acquire post-treatment data. Each animal received both psilocybin and saline, with at least 1 week between imaging sessions and the order of treatment was balanced across subjects. For calcium imaging of dendrites, 8 C57BL/6J mice including 3 males and 5 females were treated with psilocybin (244 dendritic branches including 149 from PT and 95 from IT neurons, with 4835 dendritic spines including 2637 from PT and 2198 from IT neurons) and saline (230 dendritic branches including 140 from PT and 90 from IT neurons, with 4544 dendritic spines including 2307 from PT and 2237 from IT neurons).

### Analysis of the imaging data

For structural imaging of dendrites, motion correction was performed using StackReg plug-in^9^ in ImageJ. Quantification of structural parameters such as spine head width and spine protrusion length were done according to standardized critera^10^. In brief, a dendritic spine was counted when the protrusion extended for >0.4 µm from the dendritic shaft. The line segment tool in ImageJ was utilized to measure the distances. The spine head width was determined as the width of the widest part of the spine head. Dendritic spine protrusion length referred to the distance from the tip of the head to the base at the shaft. Alterations in spine density, spine head width, and spine protrusion length were calculated as fold change compared to the value measured for each dendritic segment on the first imaging session (day −3). The raw values for spine density, spine head width, and spine protrusion length are provided in Extended Data. Spine formation rate was calculated by determining the number of newly formed dendritic spines between two consecutive imaging sessions (i.e., day −3 and day −1) divided by the total number of dendritic spines counted in the preceding imaging session (i.e., day −3). Similarly, spine elimination rate was calculated by determining the number of missing dendritic spines between two consecutive imaging sessions divided by the total number of dendritic spines counted in the preceding imaging session. To assess the longitudinal alterations in spine formation and elimination rates, we calculated the difference of the spine formation or elimination rate from the baseline rate, which was the spine formation or elimination rate for same dendritic segment before psilocybin and saline injection (between day −3 to day −1). The raw values for spine formation and elimination rates are provided in Extended Data. To divide IT neurons based on laminar position, we treated those with cell bodies residing in depth between 200 – 400 µm below the dura as layer 2/3, while those with cell bodies residing in depth between 450 µm to 650 µm as layer 5^11,12^.

For calcium imaging of dendrites, multi-page .tiff image files from one experiment were concatenated and processed with NoRMCorre^13^ in MATLAB to correct for non-rigid translational motion. As an overview, processing involved: (1) Regions of interest (ROI) corresponding to dendritic branches and spines were manually traced using an in-house graphical user interface in MATLAB; (2) The average fluorescence trace from each ROI was then processed similar to prior work^14^ to exclude background neuropil signal, and converted to fractional change in fluorescence (Δ*F*/*F*(*t*)); (3) deconvolve the fluorescence trace into discrete calcium events. Details for each of these processing steps are described below.

Dendritic branch and spine ROIs were manually traced by scrolling through the imaging frames to find putative dendritic segments (i.e., neurite segments with > 10 spiny protrusions showing a correlated pattern of fluorescence transients). First, a given branch ROI would be traced around the dendritic shaft segment using a lasso drawing tool. Next, the putative dendritic spines for that branch segment were captured using a circle drawing tool (typically 0.8 – 1.2 μm diameter ROIs). For each ROI, the pixel-wise average was calculated at each data frame to generate a fluorescence time course *F*_ROI_(*t*). Since calcium imaging was performed on the same field of view before and after drug injections, a single ROI mask was used to extract calcium signals before and after treatment. All ROI selection was done while blinded to treatment group.

Each ROI was then processed to reduce the contribution from background neuropil. Taking each ROI’s area and considering a circle with equivalent area that has radius, *r*, an ROI-specific neuropil mask was created as an annulus with inner radius 2*r* and outer radius 3*r* centered on the centroid of the ROI. Neuropil masks excluded pixels belonging to any other dendritic branch or spine ROI. To exclude neuropil mask pixels that may belong to unselected dendritic structures, we calculated the time-average signal for each pixel, taking the median amongst pixels in the mask. Pixels were excluded from the neuropil mask if their time-averaged signal was higher than the median. Finally, the remaining pixels in the neuropil mask were averaged per data frame to generate *F*_neuropil_(*t*). Each ROI had the fluorescence from its neuropil mask subtracted as follows:

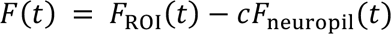

where the neuropil correction factor, *c*, was set to 0.4. Next, the fractional change in fluorescence ΔF/F(*t*) was calculated for each ROI by normalizing *F*(*t*) against its baseline, *F_0_*(*t*), estimated as the 10^th^ percentile within a two-minute sliding window:

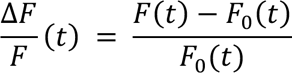

For each dendritic spine’s Δ*F*/*F*_spine_(*t*), we estimated the branch-independent spine activity, Δ*F*/*F*_synaptic_(*t*), by subtracting a scaled version of the fluorescence from the corresponding dendritic branch, Δ*F*/*F*_branch_(*t*), as follows:

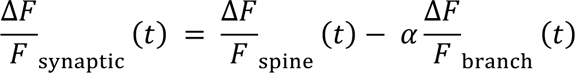

where the branch scaling factor, *α*, was computed in an ROI-specific manner using a linear regression of Δ*F*/*F*_synaptic_(*t*) predicted by Δ*F*/*F*_branch_(*t*) forced through the origin. In a previous study, we have calibration to show that with this analysis approach, the majority of the spontaneously occurring calcium transients in dendritic spines can be attributed to synaptic activation^15^.

Calcium events were detected using automated procedure for each Δ*F*/*F*_spine_(*t*) and Δ*F*/*F*_branch_(*t*) using a deconvolution “peeling” algorithm^16^. The peeling algorithm uses an iterative template-matching procedure to decompose a ΔF/F(*t*) trace into a series of elementary calcium events. The template for elementary calcium events was set to have an instantaneous onset, an amplitude of 0.3, and a single-exponential decay time constant of 1 s. Briefly, the algorithm searches a given ΔF/F(*t*) trace for a match to the template calcium event, subtracts it from the trace (i.e., “peeling”), and successively repeats the matching process until no events are found. This event detection process outputs the recorded event times with a temporal resolution by the original imaging frame rate. In this way, it is possible to detect multiple calcium events during the same imaging frame (e.g., for large amplitude transients). For each imaging session, an ROI’s calcium event rate was computed by dividing the number of calcium events by the duration of the imaging session. The calcium events were examined further by their binned amplitude (average number of calcium events per frame, among frames with at least one event detected) and frequency (number of imaging frames with at least one event, divided by the total imaging duration). The change in calcium event rate, amplitude, and frequency across treatment injections was computed for each ROI using the post-injection minus pre-injection values divided by the pre-injection values and provided raw values for calcium event rates averaged across dendritic branches in the same field of view. Separately, we have tried analyzing the Δ*F*/*F*_spine_(*t*) and Δ*F*/*F*_branch_(*t*) using a different calcium event detection algorithm OASIS^17^, which yielded qualitatively similar results (data not shown).

### Confocal imaging

After longitudinal two-photon imaging and at 3 days after psilocybin (1 mg/kg, i.p.) or saline (10 mL/kg, i.p.) injection, the mouse was deeply anesthetized with isoflurane and transcardially perfused with PBS followed by paraformaldehyde (PFA, 4% in PBS). The brains were fixed in 4% PFA for 24 hr at 4°C, and then 50-um-thick coronal brain slices were sectioned using a vibratome (VT1000S, Leica) and placed on slides with coverslip with mounting medium (Vector Laboratories #H-1500-10). The brain slices were imaged with a confocal microscope (LSM 710, Zeiss) equipped with a Plan-Apochromat 63x/1.40 N.A. oil objective (zoom 2.5) and 0.37 µm steps at 1024 × 1024 pixels with a resolution of 0.08 µm per pixel to collect the structural imaging data. In total, 204 dendrites from 10 mice were imaged for psilocybin (5 *Htr2a^f/f^* mice; 5 C57BL/6J mice), and 207 dendrites from 10 mice were imaged for saline (3 *Htr2a^f/f^* mice; 7 C57BL/6J mice).

### Overview of behavioral studies

All behavioral assays were conducted between 10:00 AM and 4:00 PM. For the animals uesd in chemogenetic manipulation, the same mice were tested on all assays. At least 2 weeks were allotted between stress-related assays. Mice were randomized into different groupings for each assay (i.e., the same mouse could be part of the psilocybin group on first assay, and then saline group on the second assay).

For studies involving PT neurons, *Fezf2-2A-CreER* mice were tested on fear extinction, then after the last extinction session 2-3 weeks later on learned helplessness, then 1-2 weeks later on head-twitch response, and finally 3 weeks later on tail suspension. We started, for fear extinction, with 58 *Fezf2-2A-CreER* mice injected with DREADD or control viruses, including 17 mice for psilocybin:mCherry (9 males, 8 females), 13 mice for saline:mCherry (7 males, 6 females), 13 mice for psilocybin:hM4DGi (8 males, 5 females), and 15 mice for saline:hM4DGi (7 males, 8 females). For learned helplessness, we had 57 *Fezf2-2A-CreER* mice remaining, including 13 mice for psilocybin:mCherry (8 males, 5 females), 15 mice for saline:mCherry (8 males, 7 females), 16 mice for psilocybin:hM4DGi (9 males, 7 females), and 13 mice for saline:hM4DGi (5 males, 8 females). For head-twitch response, we had 53 *Fezf2-2A-CreER* mice remaining, 9 mice were tested on both psilocybin and saline with 1-week interval while the rest received psilocybin or saline, including 14 mice for psilocybin:mCherry (6 males, 8 females), 13 mice for saline:mCherry (6 males, 7 females), 20 mice for psilocybin:hM4DGi (12 males, 8 females), and 15 mice for saline:hM4DGi (7 males, 8 females). For tail suspension test, we had 49 *Fezf2-2A-CreER* mice remaining, including 14 mice for psilocybin:mCherry (7 males, 7 females), 10 mice for saline:mCherry (5 males, 5 females), 13 mice for psilocybin:hM4DGi (6 males, 7 females), and 12 mice for saline:hM4DGi (8 males, 4 females).

For studies involving IT neurons, *PlexinD1-2A-CreER* mice were tested on learned helplessness, then 1-2 weeks later on head-twitch response, and finally 3 weeks later on tail suspension. We started, for learned helplessness, with 47 *PlexinD1-2A-CreER* mice injected with DREADD or control viruses, including 14 mice for psilocybin:mCherry (7 males, 7 females), 11 mice for saline:mCherry (5 males and 6 females), 11 mice for psilocybin:hM4DGi (6 males, 5 females), and 11 mice for saline:hM4DGi (5 males, 6 females). For head-twitch response, we had 47 *PlexinD1-2A-CreER* mice remaining, 4 mice were tested on both psilocybin and saline with 1-week interval while the rest received psilocybin or saline, including 15 mice for psilocybin:mCherry (8 males, 7 females), 12 mice for saline:mCherry (6 males, 6 females), 11 mice for psilocybin:hM4DGi (6 males, 5 females), and 13 mice for saline:hM4DGi (7 males, 6 females). For tail suspension test, we had 41 *PlexinD1-2A-CreER* mice remaining, including 14 mice for psilocybin:mCherry (7 males, 7 females), 9 mice for saline:mCherry (5 males, 4 females), 9 mice for psilocybin:hM4DGi (5 males, 4 females), and 9 mice for saline:hM4DGi (5 males, 4 females).

For behavioral studies involving *Htr2a^f/f^* mice, separate groups of mice were used for each behavioral test. For learned helplessness, 16 local 5-HT_2A_ receptor knockout mice were tested: 8 mice with saline (4 males, 4 females) and 8 with psilocybin (4 males, 4 females). 24 littermates injected with control virus were tested: 13 with saline (6 males, 7 females) and 11 with psilocybin (6 males, 5 females). For tail suspension test, 17 local 5-HT_2A_ receptor knockout mice were tested: 9 mice with saline (5 males, 4 females) and 8 with psilocybin (4 males, 4 females). 18 littermates injected with control virus were tested: 9 mice with saline (5 males, 4 females) and 9 with psilocybin (5 males, 4 females). For head-twitch response, 15 local 5-HT_2A_ receptor knockout mice were tested: 8 mice with saline (4 males, 4 females) and 7 mice with psilocybin (3 males, 4 females). 15 littermate controls were tested: 6 with saline (3 males, 3 females) and 9 mice with psilocybin (4 males, 5 females). For head-twitch response involving *CaMKII^Cre^*mice, 12 *CaMKIICre;Htr2a^f/f^* mice (4 males, 8 females) and 11 littermate controls (3 males, 8 females) were tested with psilocybin.

### Head-twitch response

For each mouse, deschloroclozapine (DCZ; 0.1 mg/kg, i.p.; #HY-42110, MedChemExpress) or saline (10 mL/kg, i.p.) was injected if chemogenetic manipulation was tested, and then psilocybin (1 mg/kg, i.p.) or saline (10 mL/kg, i.p.) was injected 15 min later. Head-twitch response was measured in groups of 2-3 mice, typically with psilocybin- and saline-treated mice tested simultaneously. After the injection, each mouse was immediately placed into its own plexiglass chamber (4’’ x 4’’ x 4’’), which had a transparent lid and was positioned within a sound attenuating cubicle (Med Associates). A high-speed video camera (acA1920, Basler) was mounted overhead above the chambers. We recorded videos for 10 min. Between measurements, the chambers were thoroughly cleaned with 70% ethanol. The videos were scored for head twitches by a different experimenter blinded to the experimental conditions. Previously we showed that head twitches can be quantified using magnetic ear tags^18^, however here we were concerned that the ear tag might interfere with performance in other behavioral assays so opted for video recording.

### Learned helplessness

For learned helplessness, we performed the assay using an active avoidance box with a stainless-steel grid floor and a shuttle box auto door separating the two compartments (8’’ x 8’’ x 6.29’’) inside a sound attenuating cubicle (MED-APA-D1M, Med Associates). On day 1 and day 2, there was one induction session on each day. Each session consisted of 360 inescapable foot shocks delivered at 0.2 mA for 1 – 3 s, with a random inter-trial interval ranging from 1 to 15 s. At 10-15 min after the end of the second induction session, DCZ (0.1 mg/kg, i.p.) or saline (10 mL/kg, i.p.) was given (the animals used in chemogenetic manipulation), and then psilocybin (1 mg/kg, i.p.) or saline (10 mL/kg, i.p.) was injected 15 min later. On day 3, one test session was conducted, consisting of 30 escapable foot shocks delivered at 0.2 mA for 10 s, with an inter-trial interval of 30 s. A shock would be terminated early if the mouse moved to the other compartment. Movement of the mouse was captured by beam breaks in the shuttle box. A failure was counted when the mouse failed to escape before the end of a shock. After each induction or testing session, the shuttle box was cleaned with 70% ethanol. Before each testing session, the shuttle box was cleaned with 1% acetic acid solution to provide a different olfactory context.

### Tail suspension test

Animals were tested 24 hours after administration of psilocybin (0.1 mg/kg i.p) or saline (10 mL/kg, i.p.). For chemogenetic manipulation, DCZ (0.1 mg/kg, i.p.) or saline (10 mL/kg, i.p.) was given 15 min before psilocybin or saline administration. Within a tall sound-attenuating cubicle (Med Associates), the setup included a metal bar elevated 30 cm from the floor. An animal was suspended from the metal bar by securing its tail to the bar using removable tape (NC9972972, Fisher Scientific). A small plastic tube was placed around the base of the tail to prevent tail climbing during the session. Videos of the suspended animals were recorded for 6 minutes. The behavioral apparatus was thoroughly cleaned with 70% ethanol before and after each session.

### Stress-induced resistance to fear extinction

For chronic restraint stress, we based the procedures on a published study^18^. Mice were restrained inside a cone-shaped plastic bag with openings on both ends (Decapicone, MDC200, Braintree Scientific) for 3 hours each day for 14 consecutive days. The opening corresponding to the rear of the mouse was sealed by tying a wire, leaving the mouse’s tail protruding. Restrained animals were secured in an upright position inside an empty cage and monitored frequently. At 24 hr after the end of last restraint session, we began fear conditioning and extinction procedures, which were performed using a near-infrared video fear conditioning system (MED-VFC2-SCT-M, Med Associates). Prior to each session, the mouse was brought to the behavior room for habituation for ∼30 min. The fear conditioning system was equipped with stainless-steel grid floor and was controlled by the VideoFreeze software (Med Associates). On day 1 (fear conditioning), the chamber had blank straight walls and stainless-steel grid floor. Surfaces of the chamber were cleaned with 70% ethanol (context A). Each mouse was conditioned individually in a chamber and given 3 minutes to habituate. Subsequently, it received 5 presentations of an auditory tone as the conditioned stimulus (CS; 4 kHz, 80 dB, 30 s duration). Each CS co-terminated with a footshock as the unconditioned stimulus (US; 0.8 mA, 2 s duration). A 90-s intertrial interval separated the CS + US pairings. On day 3 (fear extinction 1), for each mouse, DCZ (0.1 mg/kg, i.p.) or saline (10 mL/kg, i.p.) was injected, and then psilocybin (1 mg/kg, i.p.) or saline (10 mL/kg, i.p.) was injected 15 min later. Then 45 min later, we started test for fear extinction, while the drug is presumably still present in the brain. The chamber had two black IRT acrylic sheets inserted for a sloped roof and stainless-steel grid floor covered with a white smooth floor. Surfaces of the chamber were cleaned with 1% acetic acid (context B). Each mouse was tested individually in a chamber and given 3 minutes to habituate. Subsequently, it received 15 presentations of the CS without the US. A 15-s intertrial interval separated the CS presentations. On day 4 (retention 1), we repeated the test for fear extinction in context B. On day 17 (retention 2), we repeated the test for fear extinction in context B.

### In vivo electrophysiology

Mice were habituated to head fixation with increasing duration over several days. At least 3 hr before recording, mice were anesthetized with isoflurane and a 2-mm-diameter craniotomy was made over the medial frontal cortex (AP: 1.7 mm, ML: 0.5 mm). Cold (4°C) aCSF was used to irrigate to clear debris and reduce heating during drilling. Care was taken to minimize bleeding and keep the area clear of bone fragments. The dura was removed using a metal pin (#10130-10, Fine Science Tools). A piece of Surgifoam (#1972, Johnson & Johnson) soaked in aCSF was placed above the brain tissue, which was covered with silicon polymer (#0318, Smooth-On, Inc.) to keep the craniotomy moist and clean prior to recording. For drug administration, to avoid inserting a needle during recording session which we found to cause animal to move and therefore compromise recording stability, we used a catheter system described previously^19^. A 22-gauge intravenous catheter system (#B383323, BD Saf-T-Intima Closed IV Catheter Systems) was preloaded with psilocybin or saline and maintained at a neutral pressure. At 1 hr prior to recording, mice were briefly anesthetized with isoflurane and implanted with the intravenous catheter to their intraperitoneal cavity and the catheter was fixed with a drop of Vetbond tissue adhesive (#1469, 3M Vetbond). The mice were then head fixed and the catheter tubing was secured to the mouse holder acrylic tube with tape. Silicon polymer and Surgifoam were removed from the skull and the craniotomy was briefly irrigated with aCSF. A high-density silicon probe (#Neuropixels 1.0, IMEC) with the ground and reference shorted was coated using a 10 μL drop of CM-DiI (1 mM in ethanol; #C7000, Invitrogen). The probe was then slowly lowered (100 μm/min) into the brain using a micromanipulator (MPM; M3-LS-3.4-15-XYZ-MPM-Inverted, New Scale Technologies) to the target depth of ∼2000 μm. The probe was configured to record from 384 sites. At the target depth, we waited for the probe to settle for at least 30 min before recording began. Data were acquired using the OpenEphys software^20^ in external reference mode. Action potential and local field potentials were recorded at 30 kHz and 2.5 kHz, respectively. Once the recording began, 30 min of baseline activity was collected. The animal was then administered either psilocybin or saline via the catheter and an additional 60 min of data was collected. At the end of each recording session, optotagging was performed to identify ChR2-expressing PT or IT neurons. A fiber-coupled 473 nm laser (Obis FP 473LX, Coherent) was connected to a 200 μm optical fiber, which was mounted on the manipulator with an unjacketed end aimed at the craniotomy. The OpenEphys software was used to trigger a PulsePal (#1102, Sanworks) to drive the laser control unit to produce 20 ms pulses at 1 Hz and ∼25 mW/mm^2^ per trial. Each trial lasts for 1 s, with inter-trial interval of 980 s, and we conducted at least 500 trials.

For *Fezf2-2A-CreER* mice treated with saline, we recorded from 551 cells from 6 animals (1 male, 5 females), including 104 tagged neurons and 447 untagged other single units. For *Fezf2-2A-CreER* mice treated with psilocybin, we recorded from 572 cells from 5 animals (4 males, 1 female), including 90 tagged neurons and 482 untagged other single units. For *PlexinD1-2A-CreER* mice treated with saline, we recorded from 701 cells from 5 animals (4 males, 1 female), including 38 tagged neurons and 663 untagged other single units. For *PlexinD1-2A-CreER* mice treated with psilocybin, we recorded from 607 cells from 5 animals (4 males, 1 female), including 57 tagged neurons and 550 untagged other single units.

### Analysis of in vivo electrophysiology data

SpikeInterface^21^ was used to preprocess, spike sort, and calculate single-unit metrics. Putative single units were initially identified by Kilosort 2.5^22^ and were further manually curated in Phy (https://github.com/kwikteam/phy). Quality and waveform metrics were generated via SpikeInterface. We included units that satisfied all the following quality metrics: present for at least 90% of the recording (presence ratio), ISI violation rate less than 0.5, and amplitude cutoff of less than 0.1. To identify opto-tagged neurons, we created peri-stimulus time histograms by aligning putative single-unit spiking activity to the onset of laser stimulation. We classified opto-tagged neurons via visual inspection considering the latency to spike and reliability of spiking in response to onset of laser stimulation.

### Analysis of single-cell transcriptomics data

We accessed the “whole cortex and hippocampus 2020” SmartSeq single cell RNAseq data set made publicly available by the Allen Institute^23^. We analyzed cells that belong to neuron classes already identified by Allen Institute as layer 2/3 intratelencephalic (L2/3 IT), layer 4/5 intratelencephalic (L4/5 IT), layer 5 intratelencephalic (L5 IT), layer 6 intratelencephalic (L6 IT), layer 5 pyramidal tract (L5 PT), and layer 6 corticothalamic (L6 CT). We restricted our analyses to cells that reside in frontal cortical regions: ACA, ALM, ORB, and PL-ILA. This yielded 1403 L2/3 IT, 3118 L4/5 IT, 1541 L5 IT, 640 L6 IT, 471 L5 PT, and 2159 L6 CT neurons. We extracted expression levels for 6 genes: *Htr1a*, *Htr2a*, *Htr2b*, and *Htr2c*, which encode the 5-HT_1A_, 5-HT_2A_, 5-HT_2B_, and 5-HT_2C_ receptors, as well as *Slc17a7* and *Gad1*, which are markers for glutamatergic and GABAergic neurons. The expression level was quantified by calculating the trimmed mean (25%-75%) of log2(CPM + 1), where CPM is counts per million.

### RNA isolation and real-time PCR

Tissue around viral injection sites in the medial frontal cortex was microdissected with tools treated with RNase Away (7002, Thermo Scientific) and processed via a Dounce homogenizer in BrainBits Hibernate A (NC1787837, Fisher Scientific). Cell solution was layered on OptiPrep Density Gradient Medium (D1556, Sigma-Aldrich) and centrifuged for lipid/myelin debris removal. Cell solution was filtered and GFP+ cells were sorted (BD FACSMelody Cell Sorter). RNA was extracted from GFP+ cells using the RNeasy Plus Mini Kit (#74134, Qiagen). RNA was reverse transcribed to cDNA using the High Capacity cDNA Reverse Transcription Kit (4368814, Applied Biosystems), amplified with KAPA SYBR FAST qPCR Master Mix (KK4603, Roche) using real-time PCR (QuantStudio 7 Pro) and normalized based on reference gene *Gapdh* expression. Primers were designed with Primer3 and data were analyzed with the comparative threshold cycle method. Some steps for studies involving *CaMKII^Cre^* mice were different: tissue around the medial frontal cortex was processed via a Dounce homogenizer in TRIzol (Thermo Scientific A33251). Homogenized solution was phase separated with chloroform (Electron Microscopy Sciences 12540). RNA was precipitated with isopropyl alcohol (American Bio AB07015-0100), washed with 75% EtOH, and reconstituted in ultrapure distilled H_2_O (Invitrogen 10977-015). RNA was reverse transcribed, amplified, and analyzed as above.

### Slice electrophysiology

Brain slices were prepared as previously described^24^. Briefly, mice were first anesthetized with chloral hydrate (400 mg/kg, i.p.). After decapitation, brains were removed rapidly and placed in ice-cold (∼4°C) artificial cerebrospinal fluid (ACSF) in which sucrose (252 mM) was substituted for NaCl (sucrose-ACSF) to prevent cell swelling. Coronal slices (300 μm) were cut in sucrose-ACSF with an oscillating-blade vibratome (VT1000S, Leica). Slices were allowed to recover for ∼1-2 hr in sucrose-ACSF before commencement of recording. Slices were then placed in a submerged recording chamber in standard ACSF, and the bath temperature was raised to 32°C. The standard ACSF (pH ∼7.35) was equilibrated with 95% O_2_/5% CO_2_ and contained (in mM): 128 NaCl, 3 KCl, 2 CaCl_2_, 2 MgSO_4_, 24 NaHCO_3_, 1.25 NaH_2_PO_4_, and 10 D-glucose. Layer 5 pyramidal cells in medial frontal cortex were visualized under an Olympus BX51WI microscope using a 60X infrared objective with infrared differential interference contrast (IR/DIC) videomicroscopy. A digital CMOS camera (ORCA-spark, Hamamatsu) was used to visualize neurons in slice. Low-resistance patch pipette (3-5 MΩ) were pulled from borosilicate glass (Warner Instrument) using a horizontal micopipette puller (P-1000, Sutter Instrument). Pipettes were filled with internal solution containing (in mM): 115 K gluconate, 5 KCl, 2 MgCl_2_, 2 Mg-ATP, 2 Na_2_ATP, 10 mM Na_2_-phosphocreatine, 0.4 mM Na_2_GTP, and 10 mM HEPES, calibrated to pH 7.33. Whole-cell patch clamp recording was performed with a Multiclamp 700B amplifier (Axon Instruments). The output signal was low-pass-filtered at 3 kHz, amplified x100 and digitized at 15 kHz and acquired using Clampex 10.5/Digidata 1550A software. Series resistance, monitored throughout the experiment, was between 4-8 MΩ. Cells were discarded if series resistance rose above 8 MΩ. Liquid junction potential was not corrected. Spontaneous excitatory postsynaptic currents (sEPSCs) were recorded by clamping cells near their resting potential (≈-75 mV ± 5 mV) to minimize holding currents. After baseline recording, 20 μM 5-hydroxytryptamine creatinine sulfate (Sigma-Aldrich) was washed on for 2 min for recording in 5-HT. Subsequently, there was a washout, before 30 nM MDL100907 (Marion Merrell Dow) was washed on for 5 min, and then recordings were made in 20 μM 5-HT + 30 nM MDL100907. Analysis of spontaneous excitatory postsynaptic current (sEPSC) frequency was conducted with commercially available Mini Analysis software (Synaptosoft Inc., Decatur, GA). sEPSCs were detected and measured according to the amplitude, rise time, duration, and area under the curve (fc). Synaptic events were those with an amplitude threshold of 5 pA and area threshold of 50 fc. Traces were recorded for 60 s, and the average of 3 traces (180 s) were used for analysis for each cell/treatment.

For local 5-HT_2A_ receptor knockout mice, we recorded from 23 cells from 4 animals for the baseline, continued to obtain recording from the same 23 cells after bath application of 20 μM 5-HT, and for 7 of them recorded after bath application of 20 μM 5-HT and 100 nM MDL100907. For the *Htr2a^f/f^* mice injected with control virus, we recorded from 22 cells from 4 animals for the baseline, continued to obtain recording from the same 22 cells after bath application of 20 μM 5-HT, and for 6 of them recorded after bath application of 20 μM 5-HT and 100 nM MDL100907. Some cells did not go through all treatment conditions, because the seal had degraded and the input resistance changed significantly.

### Immunohistochemistry

Brains were sectioned using a vibratome (VT1000S, Leica) to yield 50 μm-thick coronal sections. The free-floating sections were washed 3 times with 0.3% TX-100/PBS prior to a 1 hour incubation in blocking buffer (5% normal donkey serum in 0.3% TX-100/PBS) and then incubated overnight at room temperature with anti-rabbit HTR2A antibody (1:250 dilution, #RA24288, Neuromics). Brain sections were washed with 0.3% TX-100/PBS 3 times and then incubated with secondary antibody goat anti-rabbit IgG AlexaFluor 555 (A21528, Invitrogen) at room temperature for 2 hours. Sections were washed 3 times with PBS. Sections were mounted and coverslipped with Vectashield Mounting Medium with DAPI (#H-1500-10, Vector Laboratories). Tissue sections were imaged on a Zeiss LSM 710 Confocal Microscope with a Plan-Apochromat 63x/1.40 N.A. oil objective.

### Statistics

**Supplementary Table 1** provides detailed information about the sample sizes and statistical analyses for each experiment. For behavioral studies and confocal imaging, statistical analyses were performed with GraphPad Prism 10. For two-photon imaging experiments, statistical analyses were performed based on mixed effects models using the lme4 package in R. Linear mixed effects models were used to account for repeated measures and within-subject nesting (e.g., multiple spines per branch) in a manner that makes less assumptions about underlying data than the commonly used repeated measures analysis of variance. Details about the models are described below.

For two-photon imaging of dendritic structure, analyses were performed while blind to treatment and cell type or genotype. A separate mixed effects model was constructed for each of five dependent variables related to dendritic spines: spine density, average spine head width, spine protrusion length, spine formation rate, and spine elimination rate. Each model included fixed effects terms for treatment (psilocybin vs. saline), cell type (PT vs. IT), sex (female vs. male), and time (day 1 through day 65) as factors, in addition to all second and higher-order interactions amongst these terms. The variation for repeated measures within mouse, cell, and dendrite were accounted for by including a random intercept for dendrites nested by cell nested by mice. Residuals plots were inspected visually to confirm no deviations from homoscedasticity or normality. Fixed effect *P* values were computed using likelihood ratio tests comparing the full model against a model without the effect in question. *Post hoc* two-sample *t*-tests were used to contrast psilocybin and saline groups per day, with and without splitting the sample by sex. The *P* values resulting from *post hoc t*-tests were Bonferroni-corrected for multiple comparisons. For two-photon imaging involving *Htr2a^f/f^* mice, a similar mixed effects model and *post hoc t*-tests were used, except each model included fixed effects terms for treatment (psilocybin vs. saline), genotype (*Htr2a^f/f^* vs. wild-type), and time (day 1 and day 3) as factors, in addition to all second and higher-order interactions amongst these terms. For two-photon imaging involving *Thy1^GFP^*; *Htr2a^f/f^* mice, two-factor ANOVA was used for the analyses of spine density to test the interaction between treatment (psilocybin vs. saline) and conditions (*Thy1^GFP^*; *Htr2a^+/+^*:psilocybin vs. *Thy1^GFP^*; *Htr2a^f/f^*:psilocybin vs. *Thy1^GFP^*; *Htr2a^f/f^*:saline) and time (day 1 to day 7). *Post hoc t*-tests were used to compare *Thy1^GFP^*; *Htr2a^+/+^*:psilocybin versus *Thy1^GFP^*; *Htr2a^f/f^*:psilocybin, *Thy1^GFP^*; *Htr2a^f/f^*:psilocybin versus *Thy1^GFP^*; *Htr2a^f/f^*:saline, or *Thy1^GFP^*; *Htr2a^+/+^*:psilocybin and *Thy1^GFP^*; *Htr2a^f/f^*:saline. Bonferroni correction was used for multiple comparisons.

For imaging of dendritic calcium signals, blinding procedures were implemented by having one person performed the imaging and scrambled the group names, while another person analyzed the data blind to treatment and cell type information. Data were unblinded after all the analyses were completed. A similar linear mixed effects modeling approach was used to examine three dependent variables: calcium event rate, amplitude, and frequency. Dendritic branch and spine (branch-independent) signals were analyzed in separate models (i.e., six models total). Each model included fixed effects terms for treatment (psilocybin vs. saline), cell type (PT vs. IT), and the interaction term for treatment x cell type. Treatment order (psilocybin before saline vs. psilocybin after saline) was included in the model as a nuisance variable. The variation for repeated measures of mice and dendrites were accounted for by including a random intercept for dendrites nested by field of view nested by mice. *Post hoc* two-sample *t*-tests were used to contrast psilocybin and saline groups, with and without splitting the sample by cell type. The calcium imaging statistical outputs were processed akin to the structural imaging model outputs as described above (i.e., residuals plots were inspected, fixed effect *P* values were computed with likelihood ratio tests, and *post hoc* two-sample *t*-test *P* values were Bonferroni-corrected for multiple comparisons).

For confocal imaging, two-factor ANOVA was used for the analyses of spine density, spine head width, and spine protrusion length to test the interaction between treatment (psilocybin vs. saline) and genotype (*Htr2a^f/f^* vs. wild-type). *Post hoc t*-tests were used to compare psilocybin:PT neuron-targeted 5-HT_2A_ receptor knockout versus saline: PT neuron-targeted 5-HT_2A_ receptor knockout or psilocybin:wild-type and saline:wild-type. Bonferroni correction was used for multiple comparisons.

For behavioral studies, performance was analyzed using software with automated procedures for fear extinction and learned helplessness. For head-twitch response and tail suspension test, video scoring was done by a different experimenter blinded to condition. For PT/IT studies, Two-factor ANOVA and *post hoc t*-tests were used for head-twitch response, learned helplessness test, and tail suspension test. The same statistical test was used for fear conditioning, extinction and retention to test the interaction between treatment (psilocybin vs. saline) and DREADD (hM4DGi vs. mCherry). *Post hoc t*-tests were used to compare psilocybin:mCherry versus saline:mCherry or psilocybin:hM4DGi and saline:hM4DGi for different sets of tones in a session. Bonferroni correction was applied for multiple comparisons. Due to a technical issue (faulty USB connection causing VideoFreeze software to crash in the middle of a session), a small subset of data from some mice were not used for the statistical test. For behavioral studies involving *Htr2a^f/f^* mice, two-tailed unpaired *t*-tests were used to compare control:saline versus control:psilocybin and local 5-HT_2A_ receptor knockout:saline versus local 5-HT_2A_ receptor knockout:psilocybin. For head-twitch response involving in *CaMKII^Cre^*;*Htr2a^f/f^*mice, two-tailed unpaired *t*-tests were used to compare *CaMKII^Cre^*;*Htr2a^f/f^* versus control mice. For slice electrophysiology, two-tailed unpaired *t*-tests were used to compare control:baseline versus control:+5HT, KO:baseline versus KO:+5HT, control:baseline versus control:+5HT+MDL, or KO:baseline versus KO +5HT +MDL.

For *in vivo* electrophysiology, we first identified optotagged neurons by constructing 0.1 ms bin peristimulus time histogram plots aligned to laser pulse onset. For comparison of time to first spike latency between PT and IT neurons, we conducted two-sided, two-sample Kolmogorov-Smirnov test of time to first spike for all neurons. To compare the baseline firing rates we used a two-sample, independent *t*-test. For comparing the changes in firing rates before and after administration of saline or psilocybin, we conducted a paired, two-sided *t*-test using each neuron’s baseline mean firing rate in the 30-min period before saline or drug administration (Pre) and the mean firing rate in the 60-min period after saline or drug administration (Post), and Bonferroni correction was applied for *P* values.

