## Supplementary Figures for "Pyramidal cell types and 5-HT_2A_ receptors are essential for psilocybin’s lasting drug action"

**Extended Data Figures 1–17**

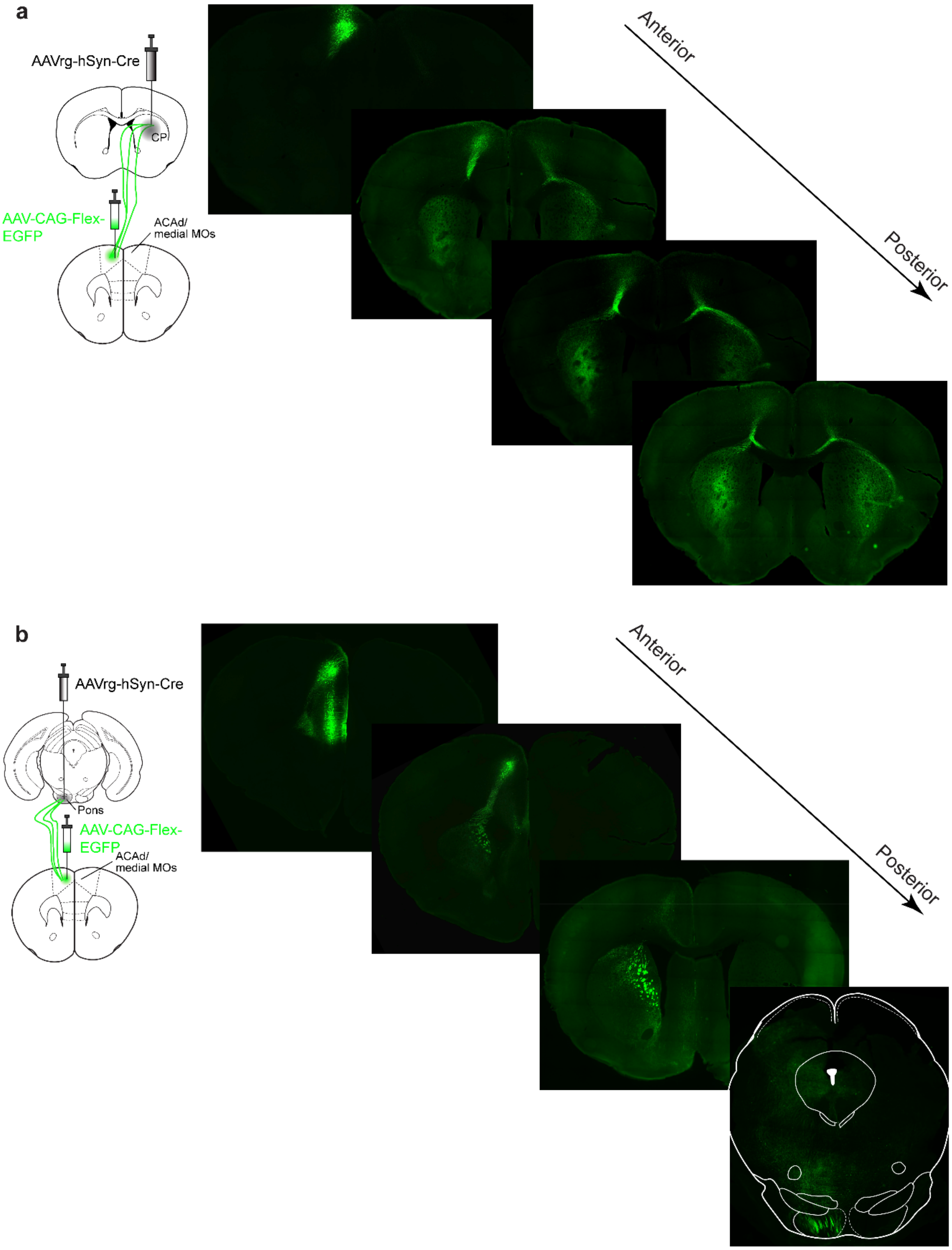


**Extended Data Fig. 1: PT and IT neurons have distinct long-range projection targets.**
**a,** To express GFP in IT neurons, we injected AAV-CAG-FLEX-EGFP in the ACAd and medial MOs portion of medial frontal cortex and low titer of the retrogradely transported AAVretro-hSyn-Cre in the contralateral striatum of adult C57BL/6J mice. Post hoc histology and imaging of the GFP fluorescence in coronal sections shows

ipsilateral and contralateral projections to various striatal and cortical regions. CP, caudoputamen. **b,** To express GFP in PT neurons, we injected AAV-CAG-FLEX-EGFP in the ACAd and medial MOs portion of medial frontal cortex and low titer of the retrogradely transported AAVretro-hSyn-Cre in the ipsilateral pons of adult C57BL/6J mice. Post hoc histology and imaging of the GFP fluorescence in coronal sections shows ipsilateral projections to striatum and subcortical regions including the pons (lower rightmost image).

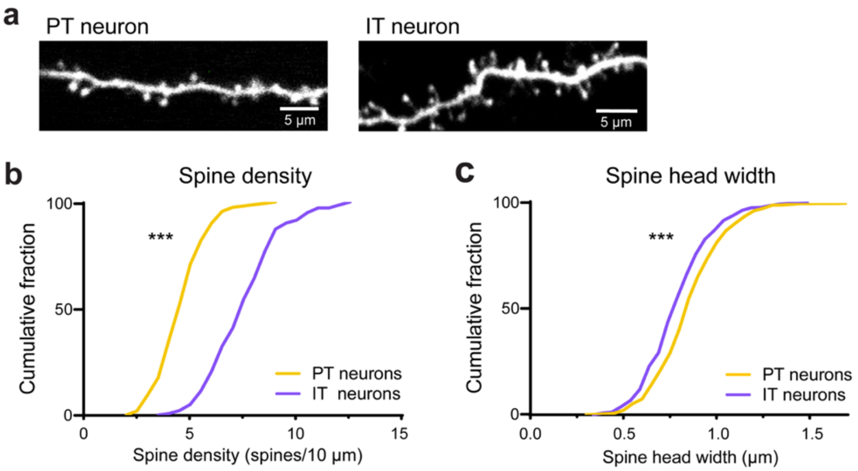


**Extended Data Fig. 2: PT neurons have lower spine density but larger spine heads than IT neurons**
**a,** *In vivo* two-photon images of apical dendrites from PT and IT neurons targeted to express GFP using retrogradely transported viruses. **b,** Baseline spine density for all imaged dendrites on Day -3 prior to any psilocybin or saline administration. PT neurons have lower spine density than IT neurons ($P < 0.001$, two-sample *t*-test). Yellow, PT neurons. Purple, IT neurons. **c,** Similar to b for spine head width. PT neurons have larger spine head width than IT neurons ($P < 0.001$, two-sample *t*-test). ***, $p < 0.001$. Sample size $n$ values are provided in Methods. Statistical analyses are provided in Supplementary Table 1.

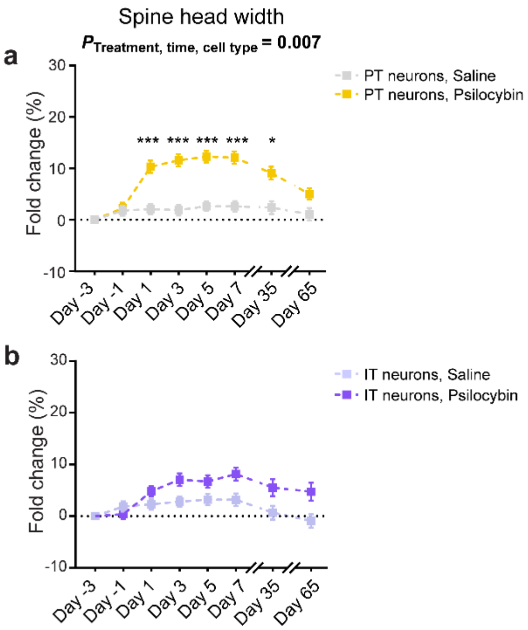


**Extended Data Fig. 3: Potential differential effect of psilocybin on dendritic spine head width in frontal cortical PT and IT neurons.**

**a,** Spine head width in the apical tuft of PT neurons after psilocybin (yellow; 1 mg/kg, i.p.) or saline (gray) across days. Mean and s.e.m. across dendrites. **b,** Similar to a for IT neurons after psilocybin (purple) or saline (light purple). There was cell-type difference in psilocybin's effect on spine head width (interaction effect of treatment × time × cell type: $P = 0.007$, mixed effects model). These results show that PT neurons have a more pronounced increase in spine head width than IT neurons, indicative of a strengthening of excitatory connections in addition to gaining new inputs for PT neurons. This enlargement in spine head for PT neurons was less durable than the changes in spine density, returning to baseline after 35 days. We note the results should be interpreted with the consideration that the spine head width is near the spatial resolution limit for *in vivo* two-photon microscopy. *, $p < 0.05$. ***, $p < 0.001$, *post hoc* with Bonferroni correction for multiple comparisons. Sample size *n* values are provided in Methods. Statistical analyses are provided in Supplementary Table 1.

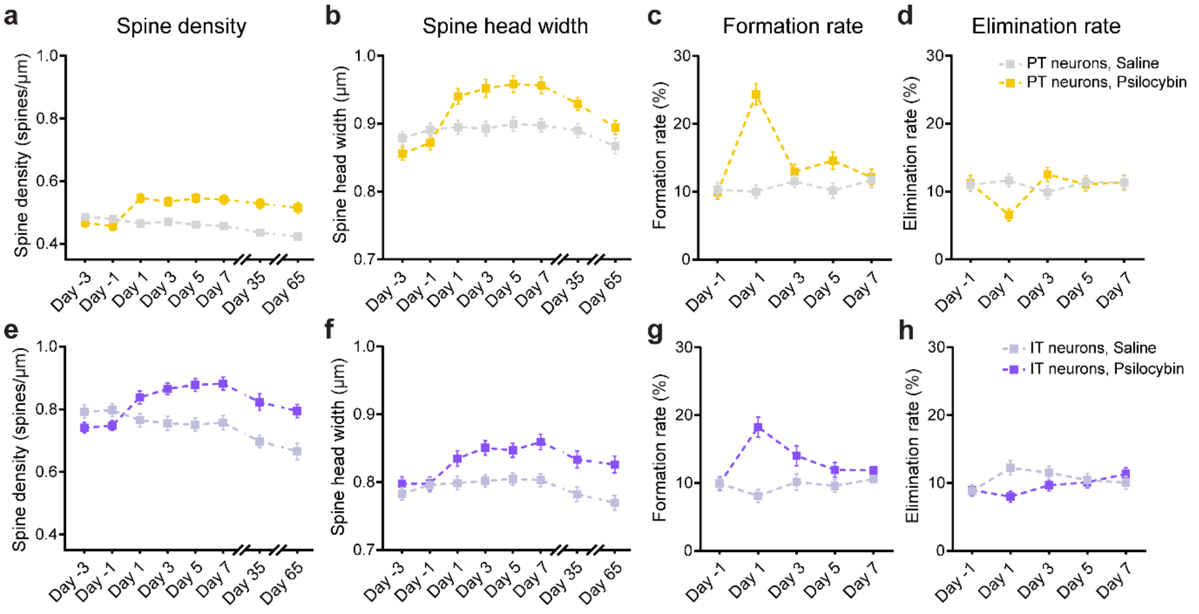


**Extended Data Fig. 4: Psilocybin induces differential structural plasticity in frontal cortical PT and IT neurons.**

**a,** Density of dendritic spines in the apical tuft of PT neurons after psilocybin (yellow; 1 mg/kg, i.p.) or saline (gray) across days. Mean and s.e.m. across dendrites. **b-d,** Similar to a for spine head width, formation rate, and elimination rate. **e-h,** Similar to a-d for IT neurons. These figure panels correspond to Fig. 1i-p, except here across-dendrite values are shown, without taking advantage of the longitudinal data for within-dendrite baseline normalization. Sample size *n* values are provided in Methods. Statistical analyses are provided in Supplementary Table 1.

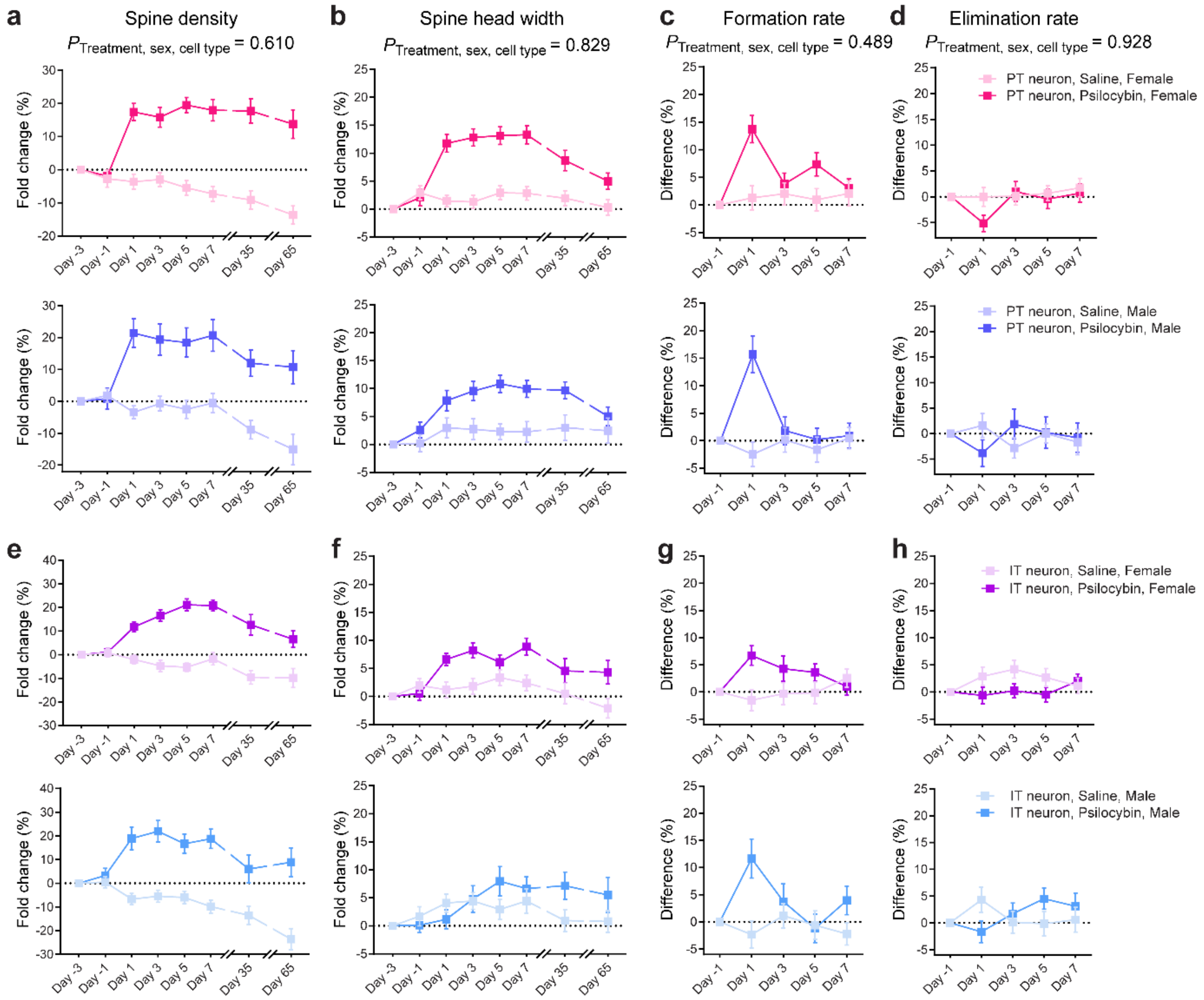


**Extended Data Fig. 5: Effects of psilocybin on structural plasticity in frontal cortical PT and IT neurons by sex of the animals.**

**a,** Density of dendritic spines in apical tuft of PT neurons in female (top row) and male mice (bottom row) after psilocybin (1 mg/kg, i.p.) or saline across days, expressed as fold-change from baseline in first imaging session (day -3). Mean and s.e.m. across dendrites. **b,** Similar to a for spine head width. **c,** Spine formation rate determined by number of new and existing spines in consecutive imaging sessions across two-day interval, expressed as difference from baseline in first interval (day -3 to day -1). **d,** Similar to c for elimination rate. **e-h,** Similar to a-d for IT neurons in female (top row) and male mice (bottom row). We did not detect effect of sex for any of the measures (interaction effect of treatment × sex × cell type, indicated in plots, mixed effects model). Sample size *n* values are provided in Methods. Statistical analyses are provided in Supplementary Table 1.

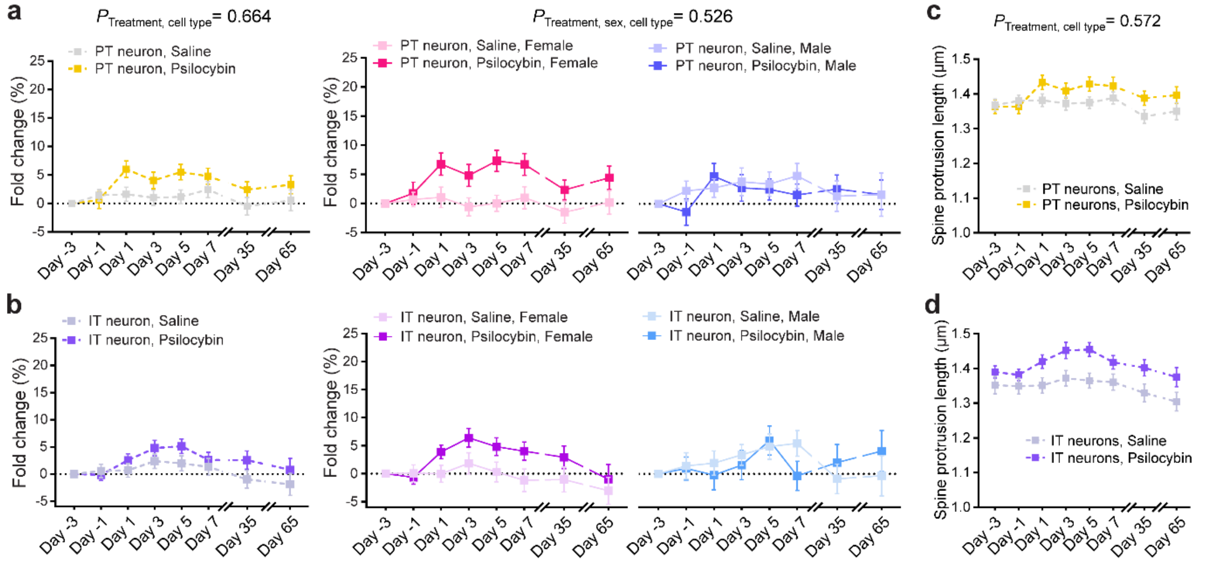


**Extended Data Fig. 6: Psilocybin has no effect on spine protrusion length.**
**a,** Protrusion length of dendritic spines in apical tuft of PT neurons for all mice (left), or separately plotted for females (middle) and males (right), after psilocybin (1 mg/kg, i.p.) or saline across days, expressed as fold-change from baseline in first imaging session (day -3). Mean and s.e.m. across dendrites. **b,** Similar to a for IT neurons. Psilocybin had no detectable effect on spine protrusion length (main effect of treatment: $P$ = 0.309, mixed effects model). **c,** Protrusion length of dendritic spines in apical tuft of PT neurons for all mice after psilocybin (1 mg/kg, i.p.) or saline across days, without taking advantage of the longitudinal data for within-dendrite baseline normalization. Mean and s.e.m. across dendrites. **d,** Similar to c for IT neurons. Sample size *n* values are provided in Methods. Statistical analyses are provided in Supplementary Table 1.

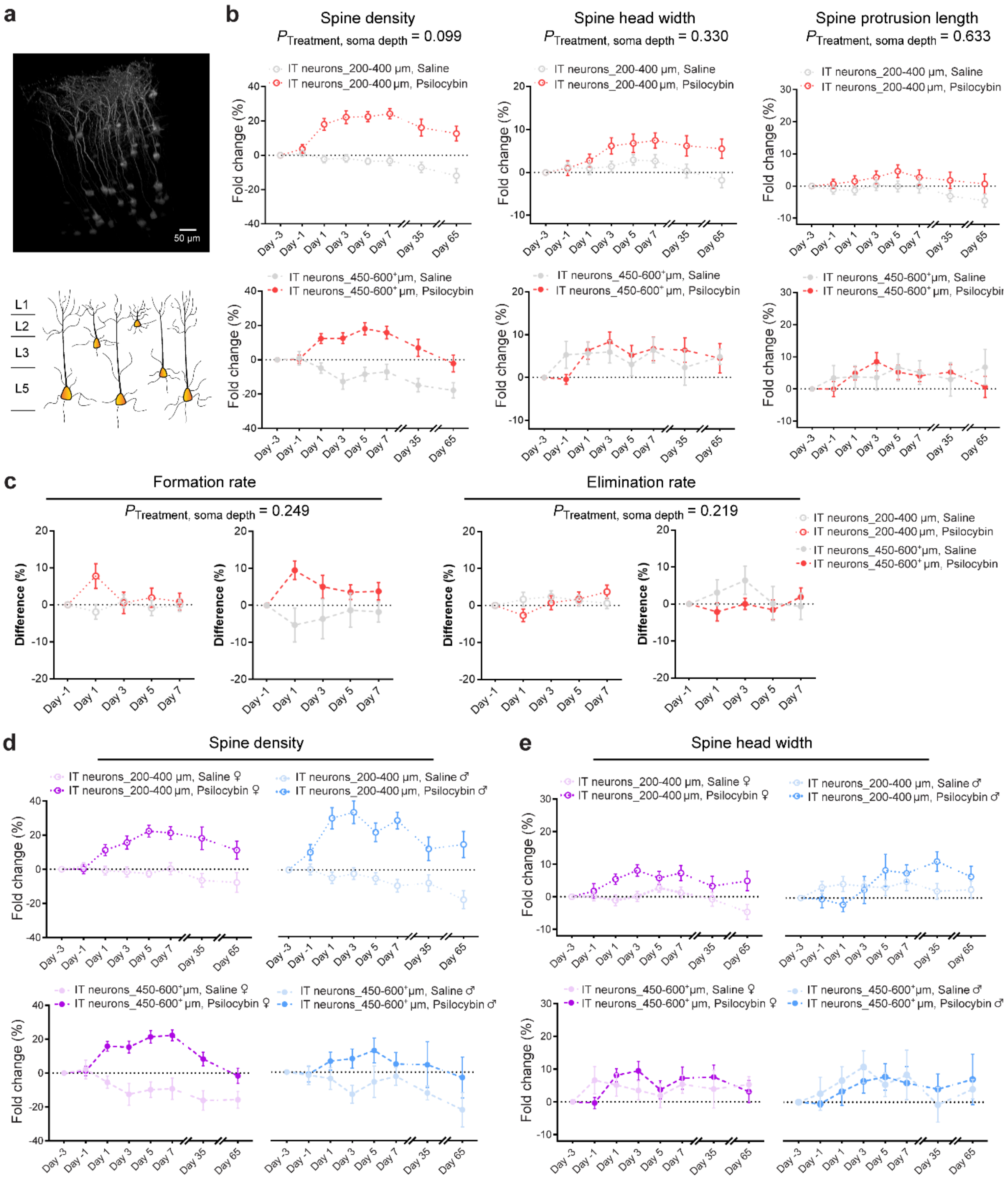


**Extended Data Fig. 7: Effects of psilocybin on structural plasticity of IT neurons residing in layer 2/3 or layer 5.**

**a,** Volumetric reconstruction from z-stack images of GFP-expressing IT neurons. IT neurons can be divided into two groups based on the laminar position of their cell body (200 – 400 µm or 450 – 600 µm). For the deep-lying IT neurons, sometimes the cell body could not be imaged due to the depth limitation of two-photon microscopy, but nonetheless the apical trunk was observed at >450 µm. **b,** Density of dendritic spines (left), spine head width (middle), and spine protrusion length (right) in apical tuft of IT neurons residing in layer 2/3 (top row) or layer 5 (bottom row) after psilocybin (1 mg/kg, i.p.) or saline across days, expressed as fold-change from baseline in first imaging session (day -3). Mean and s.e.m. across dendrites. **c,** Left: spine formation rate determined by number of new and existing spines in consecutive imaging sessions across two-day interval, expressed as difference from baseline in first interval (day -3 to day -1). Right: similar to left for elimination rate. **d-e,** Similar to b for density of dendritic spines and spine head width but further divided the data based on the sex of the animal. The analysis was motivated by the question: is psilocybin-evoked increase in spine size specific to cell type (IT versus PT), or specific to laminar position (layer 2/3 versus layer 5)? This is because IT neurons can be both superficial and deep, but PT is only found in deep layer. We detected no significant depth dependence for spine size for layer 2/3 and deep layer 5 IT neurons (interaction effect of treatment × soma depth, indicated in plots, mixed effects model). Sample size *n* values are provided in Methods. Statistical analyses are provided in Supplementary Table 1.

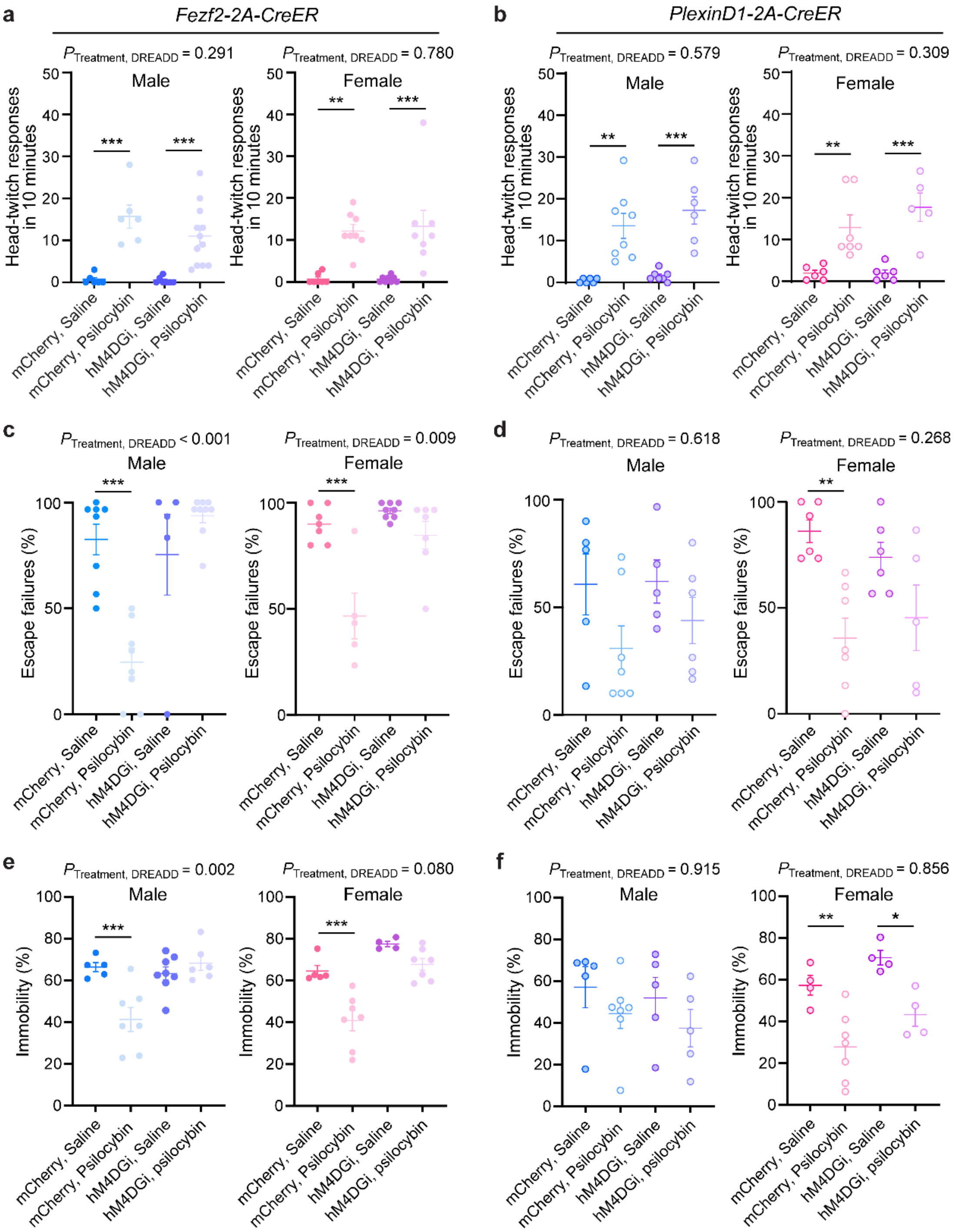


**Extended Data Fig. 8: Effects of psilocybin and chemogenetic manipulation of PT and IT neurons on behavior by sex of the animals.**

**a,** For head-twitch response assessed 10 min after drug administration, effect of chemogenetic inactivation in male (left) and female (right) Fezf2-CreER mice during psilocybin (1 mg/kg, i.p.) or saline administration. Circle, individual animal. Mean and s.e.m. **b,** Similar to a for PlexinD1-CreER mice. **c-d,** Similar to a-b for learned helplessness assessed 24 hr after drug administration. For both male and female mice, there were significant effect of PT inactivation interfering with psilocybin's impact on learned helplessness (interaction effect of treatment and DREADD: $P$ = 0.001 for males and $P$ = 0.009 for females, two-factor ANOVA). **e-f,** Similar to a-b for tail suspension test assessed 24 hr after drug administration, For male mice, there was significant effect of PT inactivation interfering with psilocybin's impact on tail suspension (interaction effect of treatment and DREADD: *P* = 0.002 for males and *P* = 0.08 for females, two-factor ANOVA). *, $p < 0.05$. ** $p < 0.01$. ***, $p < 0.001$, *post hoc* with Bonferroni correction for multiple comparisons. Sample size *n* values are provided in Methods. Statistical analyses are provided in Supplementary Table 1.

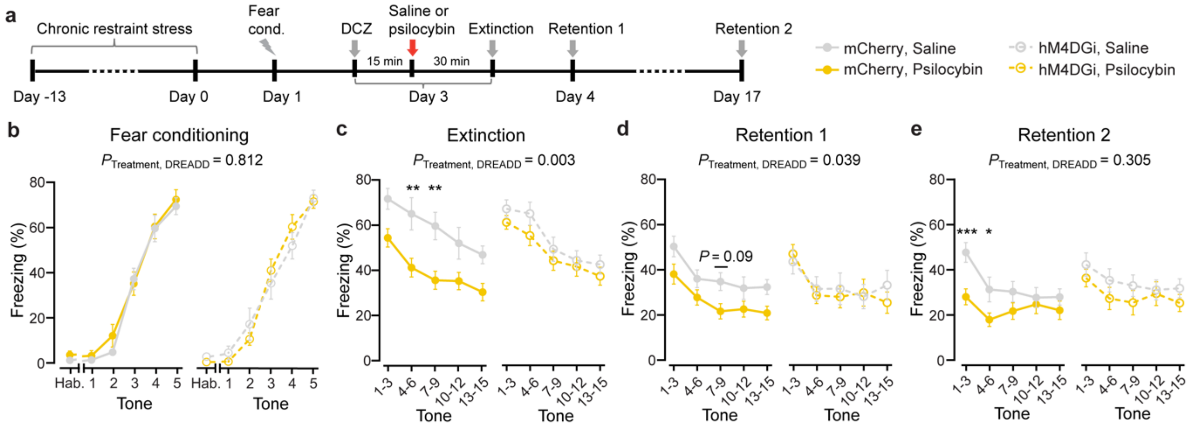


**Extended Data Fig. 9: Frontal cortical PT neurons are essential for psilocybin's facilitating effects on fear extinction.**

**a,** Chronic stress-induced resistance to fear extinction. **b,** Time spent freezing during conditioning. Mean and s.e.m. across mice. **c,** Time spent freezing during extinction session. Chemogenetic inactivation of frontal cortical PT neurons significantly diminished psilocybin's facilitating effect (interaction effect of treatment × DREADD: $P$ = 0.003, two-factor ANOVA). **d-e,** Similar to k for extinction retention sessions. *, p < 0.05. ** p < 0.01. ***, p < 0.001, *post hoc* with Bonferroni correction for multiple comparisons. Sample size $n$ values are provided in Methods. Statistical analyses are provided in Supplementary Table 1.

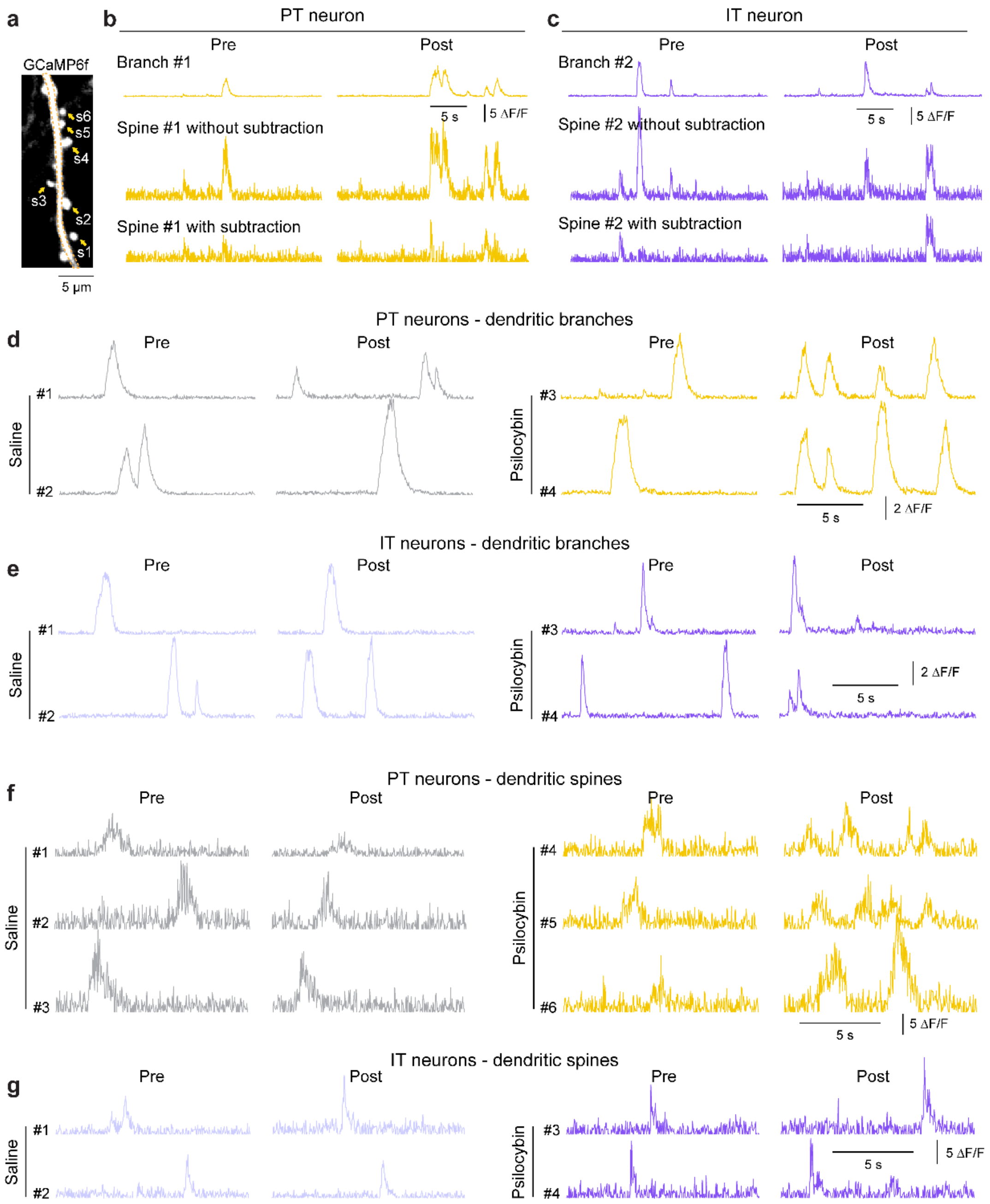


**Extended Data Fig. 10: Example $\Delta F/F_0$ traces from the dendritic branches and spines of PT and IT cells.** **a,** An example *in vivo* two-photon image of GCaMP6f-expressing apical dendrite and spines from a PT neuron. Dashed lines, the dendritic branch. Arrows, spines attached to the branch. **b,** Top, $\Delta F/F_0$ trace of a PT dendritic branch ($\Delta F/F_{\text{branch}}$). Middle, $\Delta F/F_0$ trace recorded from a dendritic spine attached to that branch ($\Delta F/F_{spine}$). Bottom, the branch-subtracted spine calcium signal ($\Delta F/F_{synaptic}$), subtracting the scaled version of $\Delta F/F_{\text{branch}}$ signal from the $\Delta F/F_{spine}$ signals. The synaptic calcium signals of dendritic spines shown and analyzed in the paper are branch-subtracted $\Delta F/F_{spine}$ transients. Left, $\Delta F/F_0$ signals before psilocybin. Right, $\Delta F/F_0$ signals after psilocybin. **c,** Similar to b for $\Delta F/F_0$ traces in an IT neuron. **d,** Left, $\Delta F/F_0$ from dendritic branches of two different PT neurons before and after saline. Right, $\Delta F/F_0$ from dendritic branches of two different PT neurons, before and after psilocybin (1 mg/kg, i.p.). **e,** Similar to d for dendritic branches of IT neuros. **f-g,** Similar to d-e for dendritic spines.

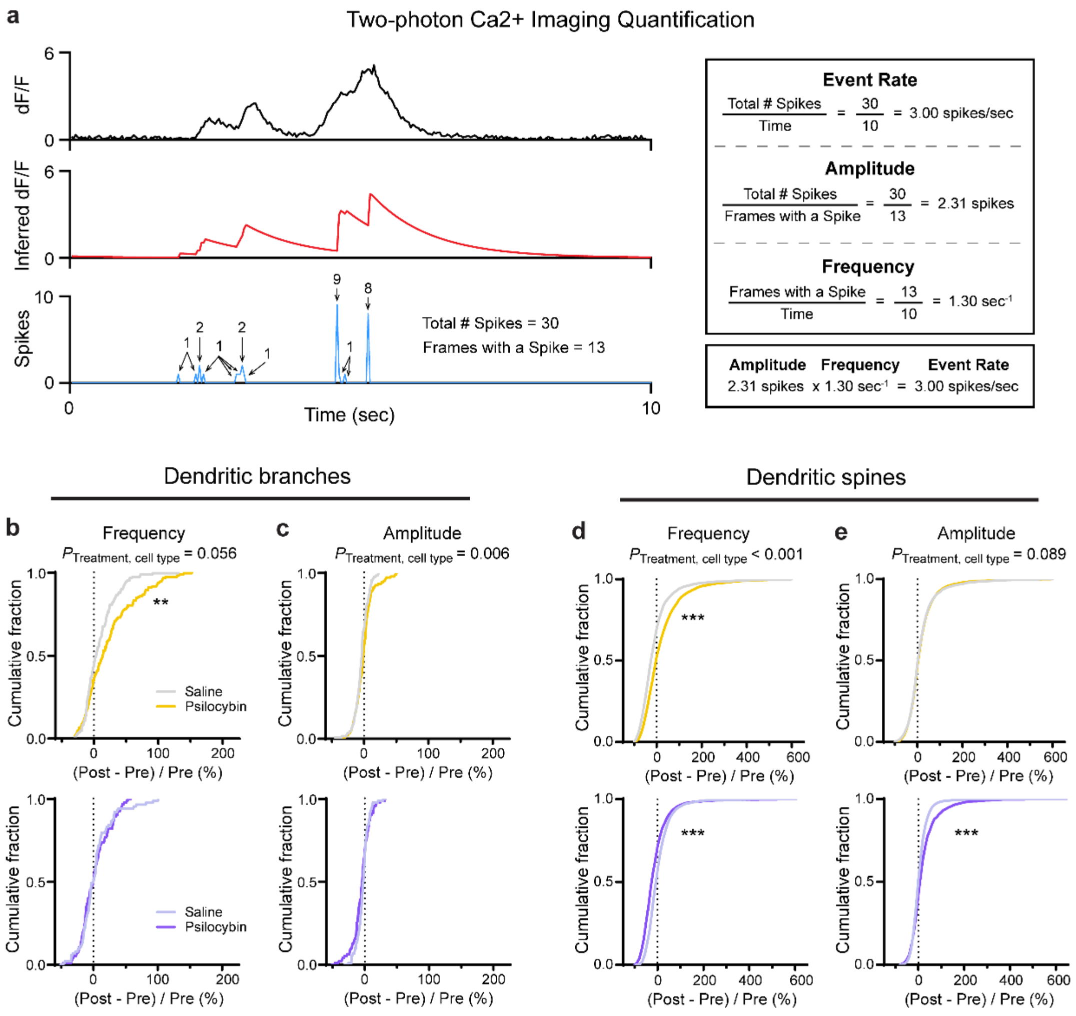


**Extended Data Fig. 11: Two-photon imaging quantification in PT and IT cells.**
**a,** Schematic illustrating how fluorescent transients are processed and analyzed to derive event rate, amplitude, and frequency. **b,** Fractional change in frequency detected in dendritic branches of PT neurons (top row) and dendritic branches of IT neurons (bottom row) after psilocybin (PT, yellow; IT, purple) or saline (PT, gray; IT, light purple). **c,** Similar to b for amplitude. **d-e,** Similar to b-c for dendritic spines. ** $p < 0.01$. ***, $p < 0.001$, *post hoc* with Bonferroni correction for multiple comparisons. Sample size *n* values are provided in Methods. Statistical analyses are provided in Supplementary Table 1.

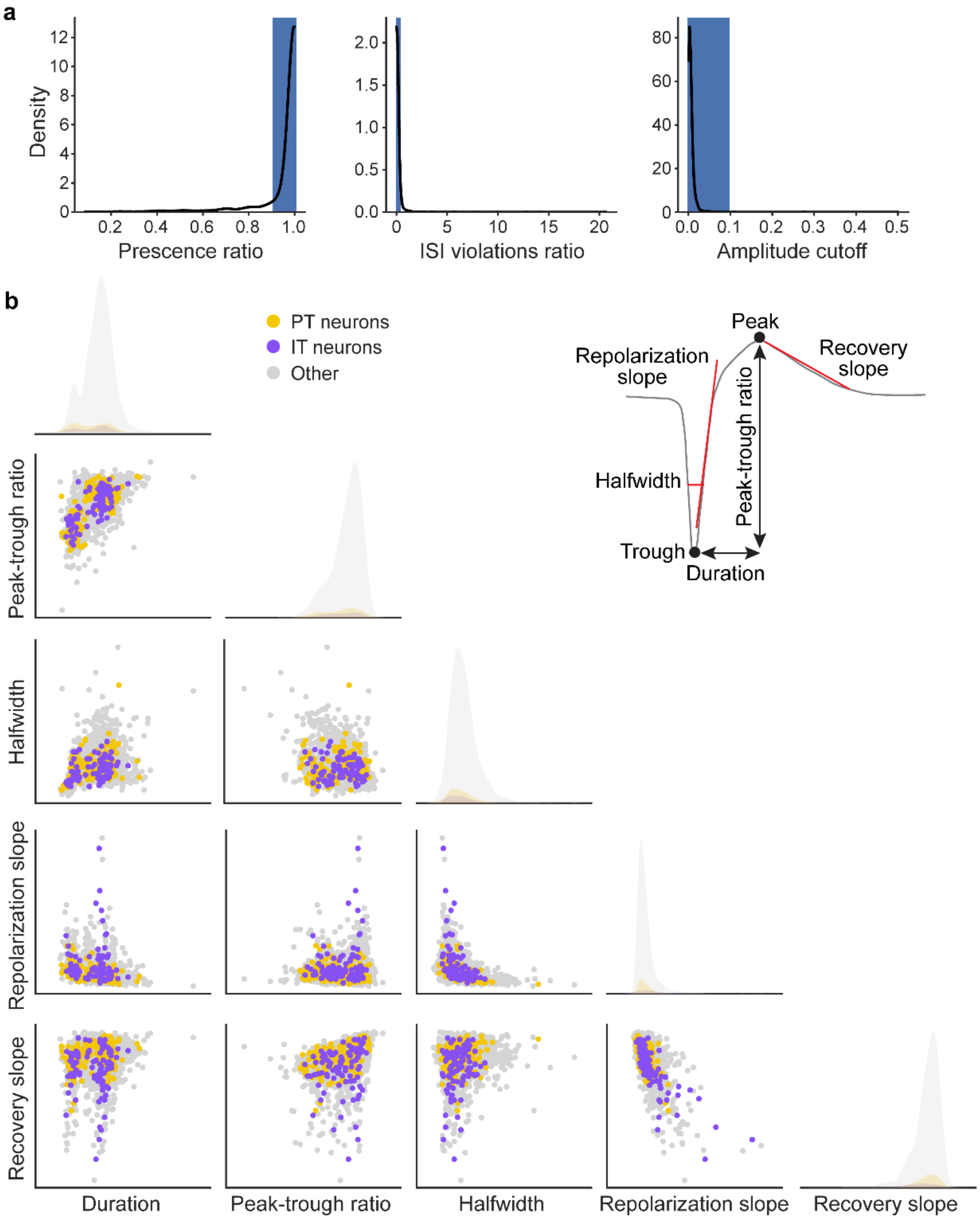


**Extended Data Fig. 12: Quality inspection and waveform analysis of recorded cells.**
**a,** The three quality metrics, including the empirical distribution of recorded units and thresholds used for curation of single units to include for the analysis of the electrophysiology data. **b,** Mean spike waveform features for all opto-tagged neurons and other untagged cells in *Fezf2-CreER* and *PlexinD1-CreER* mice. There is no single feature of the spike waveform that can reliably classify the two types of opto-tagged neurons or from the untagged cells.

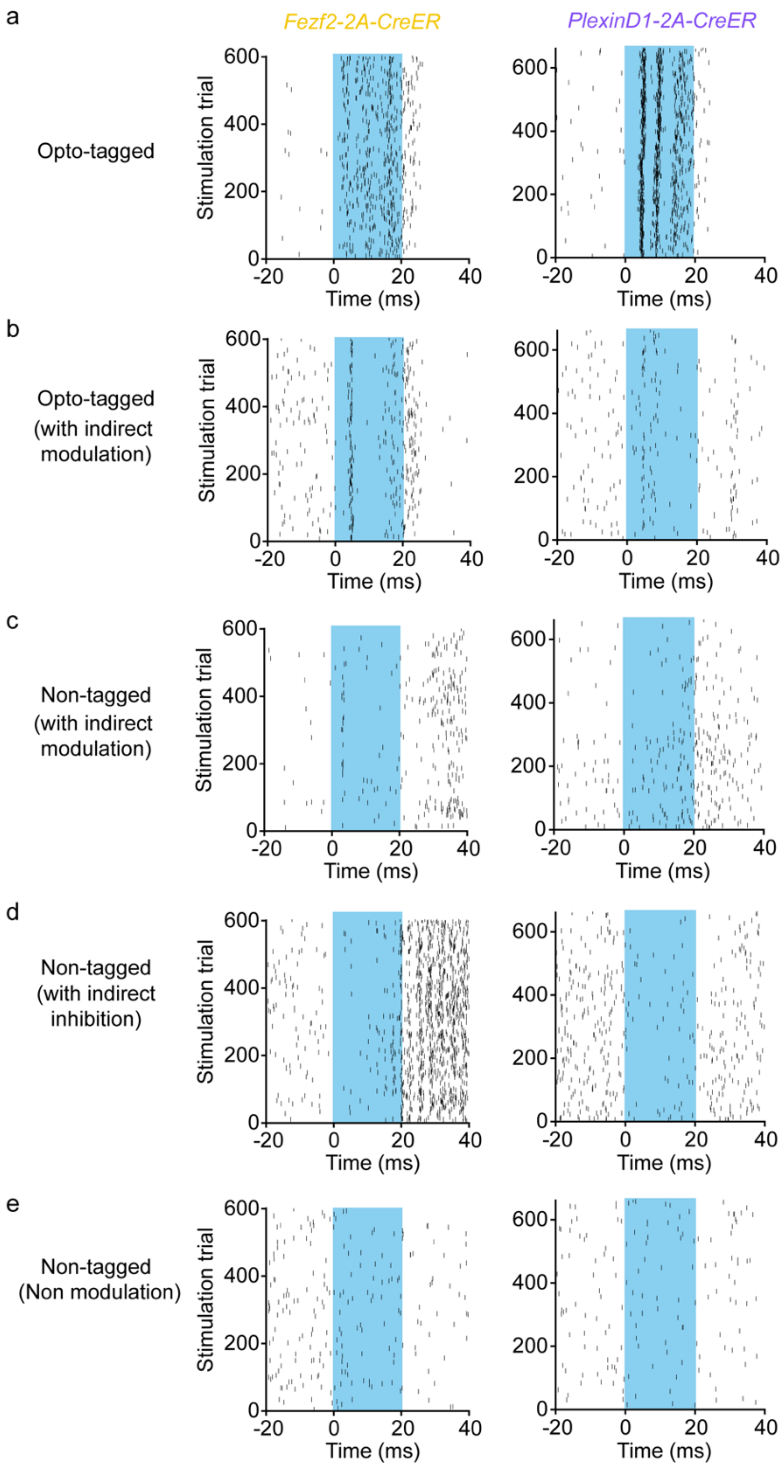


**Extended Data Fig. 13: More examples of opto-tagged neurons and untagged neurons.**
**a-b,** Spike raster of neurons classified as opto-tagged in *Fezf2-2A-CreER* (left) and *PlexinD1-2A-CreER* (right) mice. **c-e,** Spike raster of neurons classified as untagged in *Fezf2-2A-CreER* (left) and *PlexinD1-2A-CreER* (right) mice.

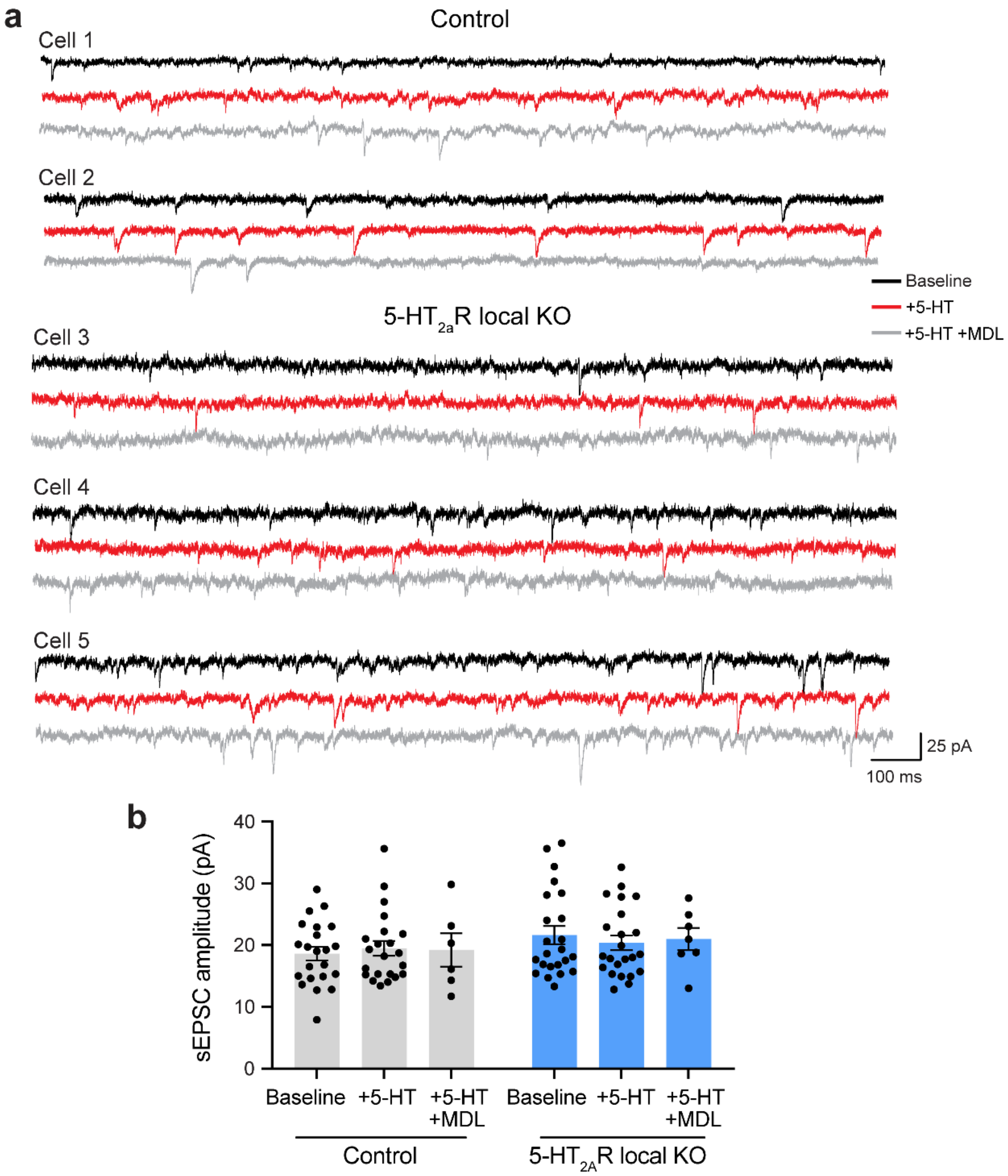


**Extended Data Fig. 14: Whole-cell voltage-clamp recordings of spontaneous EPSCs from GFP+ layer 5 pyramidal neurons**

**a,** Example sEPSC traces from 5 GFP+ layer 5 pyramidal neurons, with each set of 3 traces coming from recording of the same cell, including baseline (black), after bath application of 20 μM 5-HT (red), and after bath application of 20 μM 5-HT with 100 nM MDL100,907 (gray). Cells 1 and 2 are from control animals. Cells 3, 4, and 5 are animals with local 5-$HT_{2A}$ receptor knockout. **b,** Mean sEPSC amplitude from GFP+ layer 5 pyramidal neurons for baseline, 20 μM 5-HT, and 20 μM 5-HT + 100 nM MDL100,907 conditions, for control mice (gray) and local 5-$HT_{2A}$ receptor knockout mice (blue). Circle, individual cell. Sample size *n* values are provided in Methods. Statistical analyses are provided in Supplementary Table 1.

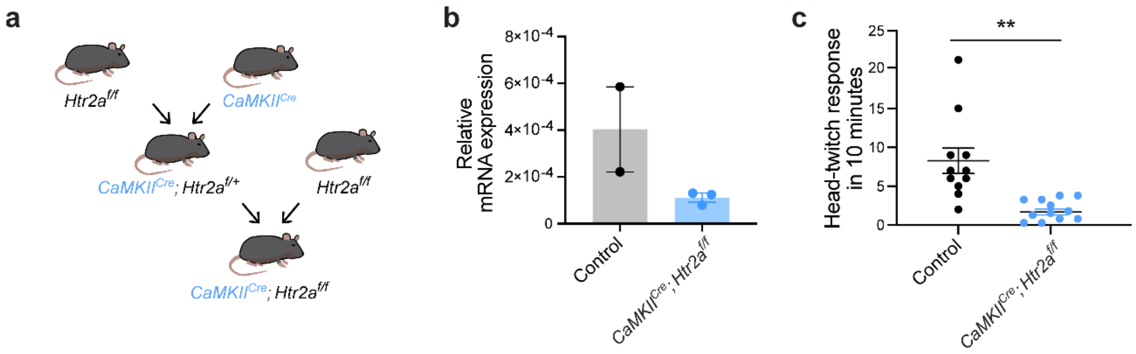


**Extended Data Fig. 15: Constitutive *CaMKII*$^{Cre}$;*Htr2a*$^{f/f}$ mice had reduced *Htr2a* transcripts and fewer psilocybin-induced head-twitch response**

**a,** Breeding scheme to generate *CaMKII*$^{Cre}$;*Htr2a*$^{f/f}$ mice. **b,** Transcript expression via qPCR from whole-brain tissue. **c,** Head-twitch response induced by psilocybin (1 mg/kg, i.p.). ** $p < 0.01$. Sample size *n* values are provided in Methods. Statistical analyses are provided in Supplementary Table 1.

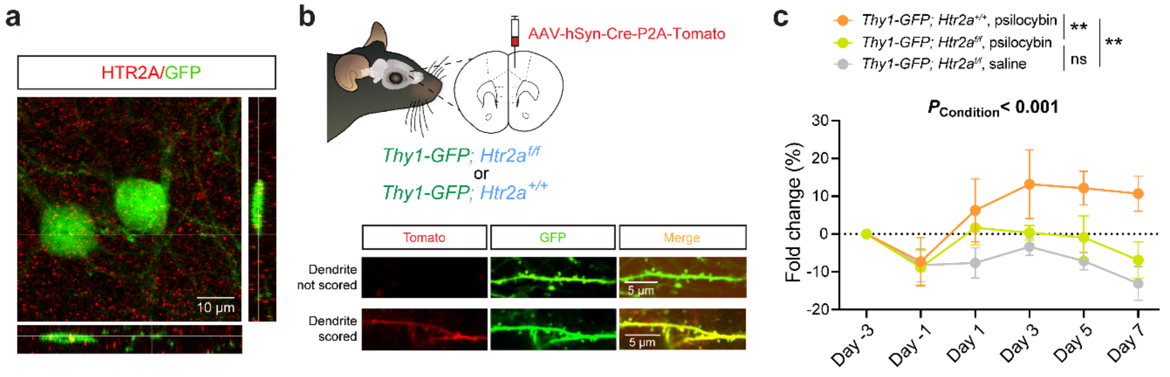


**Extended Data Fig. 16: 5-HT$_{2A}$ receptor is required for psilocybin-evoked increase of spine density in pyramidal neurons in *Thy1$^{GFP}$* mice.**

**a,** HTR2A antibody staining shows colocalization of 5-HT$_{2A}$ receptors and GFP-expressing cell bodies and neurites in the medial frontal cortex of a *Thy1$^{GFP}$* line M mouse. **b,** To image dendrites from neurons without 5-HT$_{2A}$ receptors, we injected a low titer of AAV-hSyn-Cre-P2A-Tomato into the medial frontal cortex of *Thy1$^{GFP}$*; *Htr2a$^{f/f}$* mouse. The subset of neurons with viral-mediated transgene expression would have Cre recombinase for knockout of 5-HT$_{2A}$ receptors and have tdTomato for identification. Only dendrites that expressed both tdTomato and GFP were scored. Control animals were *Thy1$^{GFP}$*; *Htr2a$^{+/+}$*. **c,** Density of dendritic spines in the apical tuft of tdTomato- and GFP-expressing neurons after psilocybin (green) or saline (gray) in *Thy1$^{GFP}$*; *Htr2a$^{f/f}$* mice or after psilocybin in *Thy1$^{GFP}$*; *Htr2a$^{+/+}$* mice (orange), expressed as fold-change from baseline in first imaging session (day -3). Mean and s.e.m. across mice. **, $p < 0.01$, *post hoc* with Bonferroni correction for multiple comparisons. Sample size *n* values are provided in Methods. Statistical analyses are provided in Supplementary Table 1.

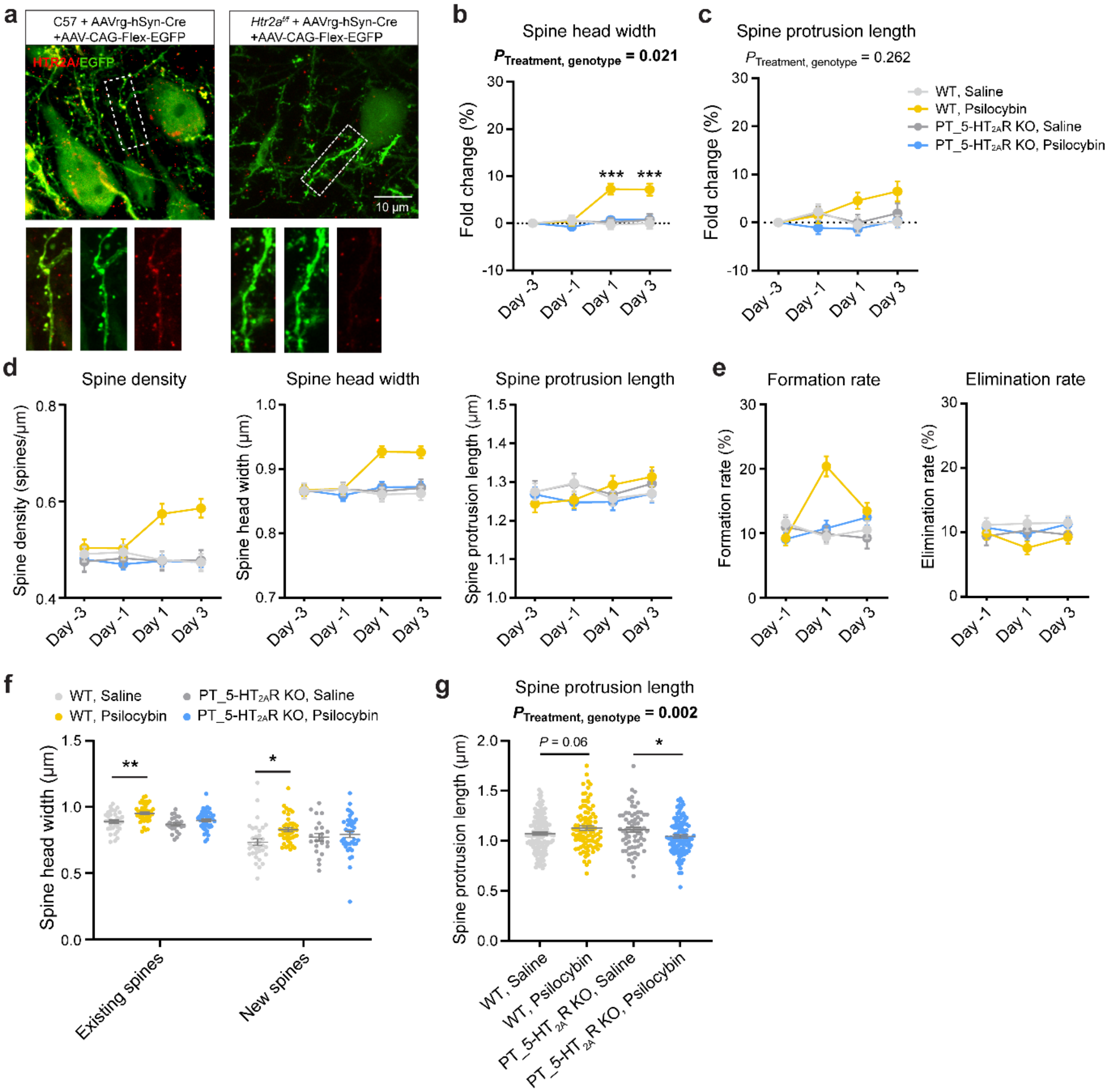


**Extended Data Fig. 17: 5-HT$_{2A}$ receptor is required for psilocybin-induced structural plasticity in PT neurons.**

**a,** HTR2A antibody staining shows colocalization of 5-HT$_{2A}$ receptors and PT neurons targeted to express GFP using retrogradely transported virus in a C57BL/6J mouse (left). For PT neuron-targeted 5-HT$_{2A}$ receptor knockout, 5-HT$_{2A}$ receptors were absent in PT neurons targeted using retrogradely transported virus in a *Htr2a*$^{f/f}$ mouse (right). **b,** From two-photon microscopy, spine head width across days, expressed as fold-change from baseline in first imaging session (day -3), in wild type mice after saline (light gray) or psilocybin (yellow) and in mice with PT neuron-targeted 5-HT$_{2A}$ receptor knockout after saline (gray) or psilocybin (blue). Mean and s.e.m. across dendrites. *Post hoc* test compared WT:saline and WT:psilocybin groups. **c,** Similar to b for spine protrusion length. **d,** Density of dendritic spines, spine head width, and spine protrusion length in the apical tuft of PT neurons across days, in wild type mice after saline (light gray) or psilocybin (yellow) and in mice with PT neuron-targeted 5-HT$_{2A}$ receptor knockout after saline (gray) or psilocybin (blue), without taking advantage of the longitudinal data for within-dendrite baseline normalization. Mean and s.e.m. across dendrites **e,** Similar to d for formation rate and elimination rate. **f,** Spine head width on day 1 plotted separately for pre-existing versus newly formed spines in different conditions. **g,** From confocal microscopy, protrusion length of dendritic spines in the apical tuft of PT neurons in wild type mice after saline (light gray) or psilocybin (yellow) and in mice with PT neuron-targeted 5-$HT_{2A}$ receptor knockout after saline (gray) or psilocybin (blue). Circle, individual dendritic segment. Mean and s.e.m. across dendrites. * $p < 0.05$. **, $p < 0.01$. ***, $p < 0.001$, *post hoc* with Bonferroni correction for multiple comparisons. Sample size *n* values are provided in Methods. Statistical analyses are provided in Supplementary Table 1.
